# The Actin Bundling Protein Fascin Promotes Lung Cancer Growth and Metastasis by Enhancing Glycolysis and PFKFB3 Expression

**DOI:** 10.1101/2021.04.02.437070

**Authors:** Shengchen Lin, Yunzhan Li, Dezhen Wang, Chongbiao Huang, David Marino, Oana Bollt, Chaodong Wu, Matthew Taylor, Wei Li, Gina M. DeNicola, Jihui Hao, Pankaj K. Singh, Shengyu Yang

## Abstract

Fascin is a pro-metastasis actin bundling protein upregulated in essentially all the metastatic carcinoma. It is believed that fascin promotes cancer cell migration and invasion by facilitating membrane protrusions such as filopodia and invadopodia. Aerobic glycolysis is a key feature of cancer metabolism that provides critical intermediate metabolites for tumor growth and cell proliferation. Here we report that fascin increase glycolysis in lung cancer to promote tumor growth and metastasis. Fascin promotes glycolytic flux by increasing the expression and activities of phosphofructose kinase 1 and 2 (PFK1 and 2). The glycolytic function of fascin depends on activation of YAP1 through its canonical actin bundling activity. Fascin promotes the binding of YAP1 to a TEAD1/4 binding motif located 30 bp upstream of the PFKFB3 transcription start site to activate its transcription. Our interrogation of the TCGA database suggest that the fascin-YAP1-PFKFB3 circuit is likely conserved across different types of cancer. We further showed that the glycolytic function of fascin is essential for promotion of lung cancer growth and metastasis. Importantly, pharmacological inhibitors of fascin could be used to suppress the YAP1-PFKFB3 signaling and inhibit glycolysis in cancer cell lines, organoid cultures and xenograft metastasis models. Taken together, our data reveal an important glycolytic role of fascin in lung cancer metabolism, and suggest that pharmacological inhibitors of fascin could be used to reprogram cancer metabolism in lung cancer and potentially other cancer with fascin upregulation.

## Introduction

The actin cytoskeleton in metastatic cancer cells is frequently dysregulated to promote cell migration, invasion and metastatic dissemination. Fascin is a monomeric actin bundling protein upregulated in essentially all the metastatic carcinoma (1–3). Fascin crosslinks actin filaments into straight and stiff bundles through its positively charged actin-binding surfaces (1, 4, 5). In addition to its well-known role in metastatic dissemination, fascin also increases metabolic stress resistance, chemoresistance and cancer cell stemness (6–8). Mechanisms underlying these non-canonical functions of fascin are not well understood.

Metastasis is responsible for more than 90% of cancer related death and there are few effective therapies for metastatic cancer (9). Fascin has emerged as an attractive target for the development of anti-metastasis therapeutics (7, 10–13) for several reasons. First, in adult animals fascin expression is absent in most tissues with the exception of neuronal tissues (1–3). Second, fascin expression is highly upregulated in transformed cancer cells, especially in metastatic cancer and fascin plays a causative role in promoting cancer metastasis (1, 2). Third, adult mice with homozygous deletion of fascin appeared to be healthy and fertile (14), which suggested that inhibitors specifically targeting fascin would likely be well tolerated in cancer patients. Several small molecules, including migrastatine analogs and G2 derivatives, have been identified as fascin inhibitors through chemical biology approaches and high throughput screening (12, 15, 16). Some FDA-approved drugs also showed inhibitory activities toward fascin (17). These small molecules were able to inhibit cell migration and invasion in cell culture and block metastatic dissemination in animal models. However, the usefulness to block metastatic dissemination alone in the clinic has been questioned (1, 9), since most cancer patients with metastatic disease have already disseminated cancer cells at the time of diagnosis.

Cancer cells undergo profound metabolic reprogramming during tumor initiation and progression (18). In metastatic cancer cells, metabolic reprogramming is essential for the colonization of distant organs, since distant sites likely have drastically different nutrient and metabolic requirements than the primary tumor (18). It is well documented that cancer cells prefer to use glycolysis for their glucose catabolism even under oxygen replete conditions (the Warburg effect) (19). Although this phenomenon was initially thought to be due to defective mitochondria in cancer, mitochondrial respiration is actually functional in most cancer cells; both OXPHOS and glycolysis are essential for tumor growth (19). The glycolytic metabolism provides crucial intermediate metabolites for the synthesis of nucleotides, amino acids and lipids, which are building blocks required for the growth and proliferation of cancer cells (19). Several glycolytic enzymes such as aldolase are actin binding proteins and their activities could be regulated by the actin cytoskeleton remodeling (20). However, the metabolic function of the actin cytoskeleton dysregulation in metastatic cancer cells remained poorly elucidated.

Targeting the deregulated cancer metabolism has emerged as an attractive new approach in the development of cancer therapeutics. A challenge in targeting specific metabolic pathway is the intrinsic metabolic flexibility, where cancer cells are able to alternate between glycolysis and mitochondrial OXPHOS in response to metabolic stress (21). Therapeutic modalities simultaneously targeting glycolysis and mitochondrial OXPHOS have improved efficacies (21). We previously reported that fascin is able to augment mitochondrial oxidative phosphorylation in lung cancer by remodeling mitochondrial actin filament and promoting biogenesis of respiratory Complex I (7). Here we show that fascin upregulates phosphofructose kinases (PFK) to promote glycolysis in lung cancer. Our data here, together with our earlier findings, suggested that fascin could be targeted to suppress both glycolysis and mitochondrial OXPHOS. Therefore, pharmacological inhibitors of fascin could be useful to preventing lung cancer metastasis by reducing metabolic flexibility in lung cancer.

## Results

### Fascin promotes glycolysis in NSCLC cells through its actin crosslinking activity

We have previously reported that knockdown (KD) or knockout (KO) of fascin in NSCLC inhibited OXPHOS (7). In response to inhibition of OXPHOS, cells usually increased glycolytic metabolism as a compensatory response (21, 22). However, we noted that fascin KO in NSCLC cells reduced both lactate production and glucose uptake by 20-30% (Fig. 1A-1B, Fig. S1A- S1B), which suggested inhibition of glycolysis in these cells. Conversely, ectopic expression of fascin in these cells increased both lactate production and glucose uptake (Fig. 1A-1B, Fig. S1A- S1B). The measurement of extracellular acidification rate (ECAR) confirmed that fascin KD or KO decreased ECAR (Fig. 1C-1D and Fig. S1C), while fascin OE increased ECAR (Fig. 1E-1F and Fig. S1D), which suggested that fascin activates glycolysis in addition to its previously reported function in augmenting mitochondrial OXPHOS.

**Figure 1.**
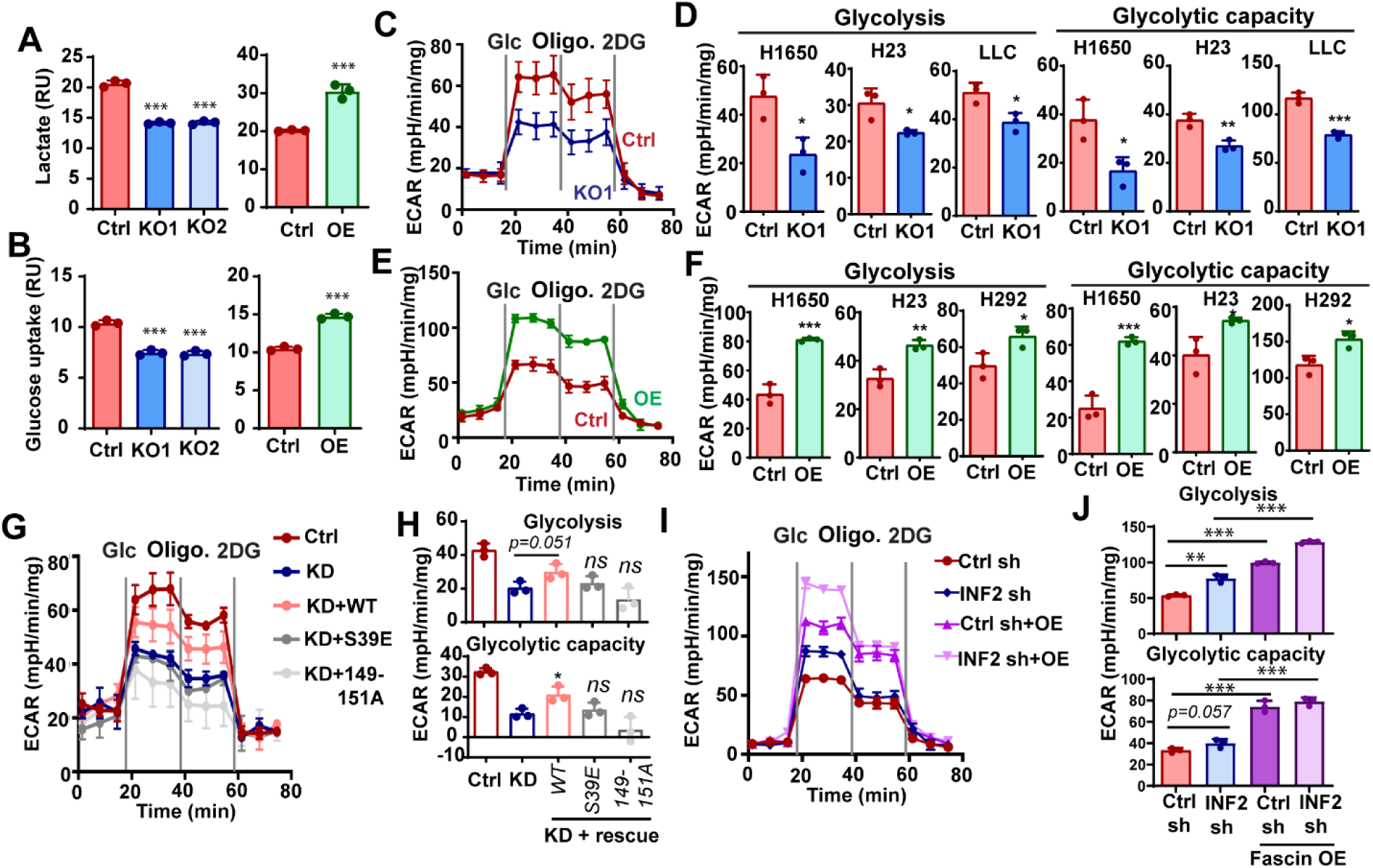
Fascin promotes glycolysis in NSCLC cells through its actin crosslinking activity. A and B, the effect of fascin KO or OE on lactate production (A) or glucose uptake in H1650 cells. C and D, the effects of fascin KO (C) or OE (D) on extracellular acidification rate (ECAR), as determined by glycolysis stress test in H1650 cells. E and F, quantitation of the effects of fascin KO (E) or OE (F) on glycolysis and glycolytic capacity in different NSCLC cell lines. G and H, the effect of fascin knockdown (KD) or rescue with ectopic expression of wild-type (WT) fascin or bundling defective mutants (S39E and 149-151A) on ECAR in H1650 cells. The ectopic WT fascin, but not the bundling defective mutants, was able to rescue ECAR, including glycolysis and glycolytic capacity, in fascin KD cells. H is the quantitation of glycolysis and glycolytic capacity for G. I and J, the effect of fascin overexpression on ECAR in H1650 cells stably expressing control shRNA (Ctrl sh) or INF2-CAAX shRNA (INF2 sh). The ectopic expression of wild-type fascin in INF2-CAAX knockdown H1650 cells was able to further increase the ECAR, including glycolysis and glycolytic capacity (J). Data in A, B, E, F, H and J were analyzed by two-tailed, two-sample unpaired Student’s test. Results from at least three independent experiments are shown. *, ** and **** indicated p<0.05, 0.01 and 0.001, respectively. ns, not significant.

Next, we asked whether fascin regulates NSCLC glycolysis through its canonical actin bundling activity. We have previously demonstrated that mutations at the two positively-charged actin binding surfaces were able to abrogate fascin crosslinking activities without causing significant conformation change (4, 23). Therefore we ectopically expressed wild type fascin (WT) or two bundling-defective mutants (S39E and 149-151A) in fascin knockdown cells. As shown in Fig. 1G and 1H, the ectopic expression of WT fascin, but not the bundling defective mutants, was able to at least partially restore ECAR in fascin KD cells. Since fascin controls mitochondrial OXPHOS by remodeling mitochondrial actin filaments (mtF-actin), also in an actin-bundling activity dependent manner, we next asked whether mtF-actin might be involved in fascin-mediated glycolysis. We disrupted mtF-actin by knocking down INF2-CAAX, the endoplasmic membrane actin nucleator involved in the polymerization of mtF-actin (24). INF2-CAAX knockdown abrogates the accumulation of mtF-actin and fascin-mediated enhancement of mitochondrial OXPHOS in lung cancer cells (7). As shown in Fig. 1I-1J, INF2-CAAX knockdown modestly increased ECAR in H1650 cells, likely due to the compensatory response to the decrease in mitochondrial OXPHOS in these cells (7). The overexpression of fascin in INF2-CAAX knockdown cells was able to increase ECAR to an extent similar to that in control shRNA cells (Fig. 1I-1J), which suggested that mtF-actin is not required for fascin to promote glycolysis. Taken together, our data support that fascin is able to promote glycolysis in addition to its previously reported role in augmenting mitochondrial OXPHOS. The glycolytic function of fascin requires its actin bundling activity but is independent of mtF-actin remodeling.

### Fascin enhances the glycolytic flux to fructose-1, 6-bisphosphate by upregulating PFKs

To understand the mechanism by which fascin regulates glycolysis, we used ^13^C-labeled glucose to investigate the effect of fascin KO on glycolytic flux in H1650 cells. Cells were incubated with U-^13^C-Glucose for 0, 2, 15, 30, 60 or 360 minutes and the incorporation of ^13^C to several glycolytic intermediate metabolites were analyzed using LC-MS/MS. As shown in Fig. 2A, the levels of ^13^C-labeled Glucose/Fructose-6-phosphate (G6P /F6P), 2 / 3-phosphoglyceric acid (2PG / 3PG), Phosphoenolpyruvic acid (PEP) and Fructose bisphosphate (FBP) were mostly lower in fascin KO cells when compared to control H1650 cells in earlier time points (2-60 minutes) (Fig. 2A). After 360 minutes most of the ^13^C-labeled glycolysis metabolites in KO cells returned to levels comparable to (or in the case of G6P/F6P higher than) control cells except for FBP (Fig. 2A). FBP was markedly lower in KO cells even at 6 hours. Fructose-1,6-bisphosphate (F-1,6-BP) is a product of the phosphorylation of F6P by PFK1. Along with hexokinase (HK) and pyruvate kinase (PK), PFK1 catalyze one of the three irreversible and rate limiting steps in the glycolysis cascade (19). The activity of PFK1 is allosterically regulated by Fructose-2,6- bisphosphate, a product of PFK2. We confirmed that fascin KO decreased, while fascin OE increased, the levels of Fructose-1, 6-bisphosphate in H1650 and H292 cells (Fig. 2B and 2C). Fructose-1, 6-bisphosphate allosterically activates PKM2 (Fig. 2A, left panel), the pyruvate kinase isoform upregulated in cancer, by promoting its tetramerization (25). As shown in Fig. 2D and 2E, fascin KO or KD decreased the levels of PKM2 tetramers and dimers while increasing the levels of the monomeric protein. The ectopic expression of WT fascin, but not the bundling defective mutants, was able to restore the levels of PKM2 tetramer levels in fascin KD cells. These data are consistent with the notion that fascin KO reduced the glycolytic flux to F-1,6-BP.

**Figure 2.**
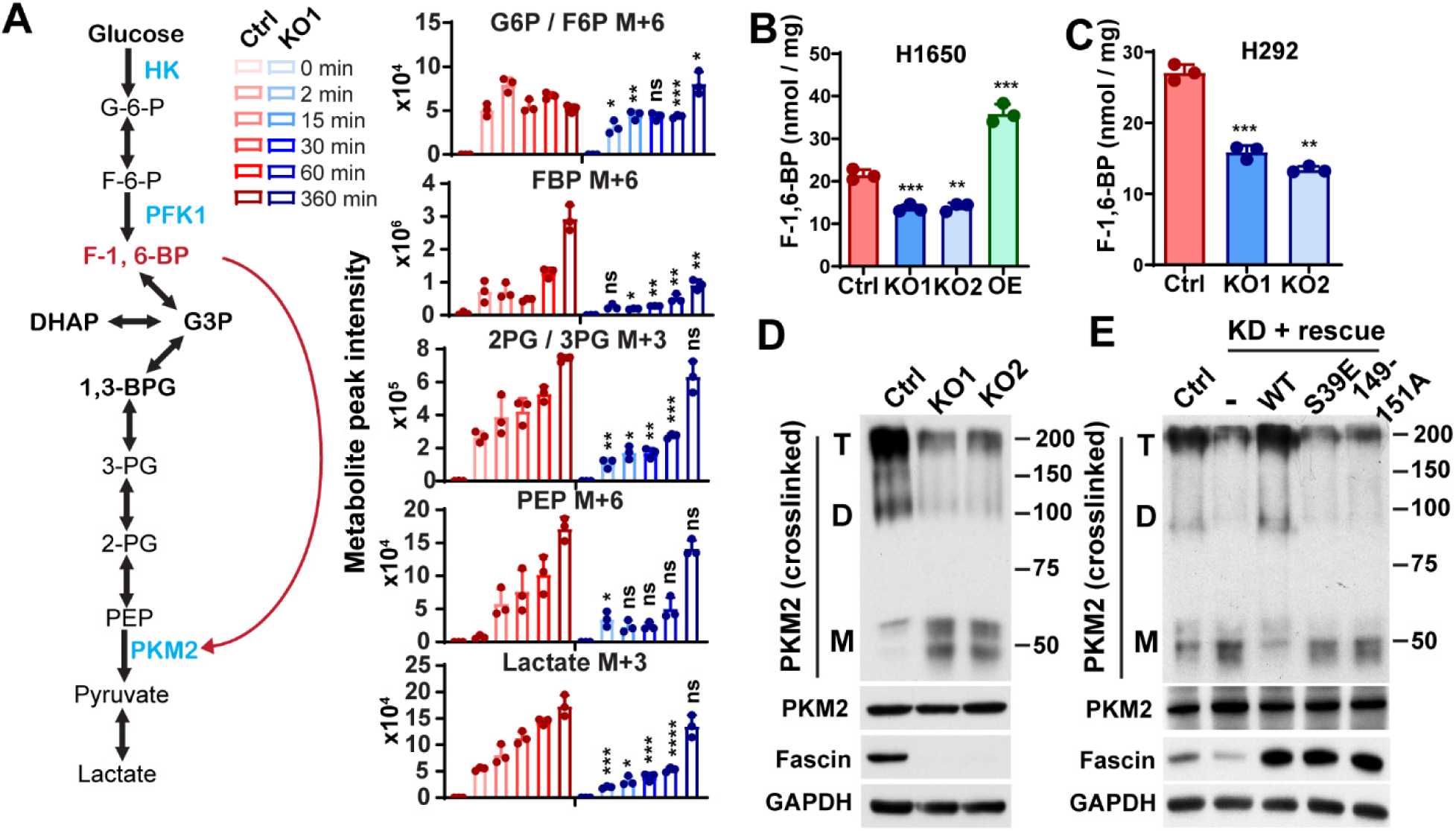
Fascin enhances the glycolytic flux to fructose-1, 6-bisphosphate. A, left panel, the schematic illustration of glycolysis pathways. The red arrow indicates F-1,6-BP can bind to PKM2 to activate its tetramerization and catalytic activity. Right panel, the relative level of ^13^C-labeled glycolytic metabolites in control or fascin KO H1650 cells after incubation with U-^13^C-Glucose for indicated time. M+x indicates the number of ^13^C-labeled carbon in respective metabolites. B and C, the effect of fascin KO on F-1,6-BP levels in H1650 and H292 cells. D, Western blotting showing that fascin KO reduced the levels of PKM2 dimer and tetramer without affecting the expression of total PKM2 protein levels in H1650 cells. T, D, M indicated tetramer, dimer and monomer. E, the ectopic expression of wild-type fascin, but not the bundling defective mutants (S39E and 149-151A), in fascin knockdown H1650 cells was able to rescue PKM2 dimer and tetramer level in H1650 cells. T, D, M indicated tetramer, dimer and monomer. Data in A- C were analyzed by two-tailed, two-sample unpaired Student’s test. Representative results from at least three (B-E) or two (A) independent experiments are shown. *, **, *** and **** indicated p<0.05, 0.01, 0.001 and 0.0001, respectively. ns, not significant.

To understand the mechanism by which fascin regulates FBP production, we examine the effect of fascin KO on the expression levels of three isoforms of PFK1 (PFKP, PFKM and PFML) and PFKFB3, the PFK2 isoform frequently upregulated in cancer. As shown in Fig. 3A, fascin KO significantly reduced the protein levels of PFKP, PFKM and PFKFB3, but not that of PFKL. Consistent with the reduction in PFK protein levels, the PFK activity in NSCLC cells were also lower in fascin KO NSCLC cells (Fig. 3B). Conversely, the ectopic expression of fascin was able to increase the protein levels of PFKFB3, PFKP and PFKM, but not PFKL (Fig. 3C).

**Figure 3.**
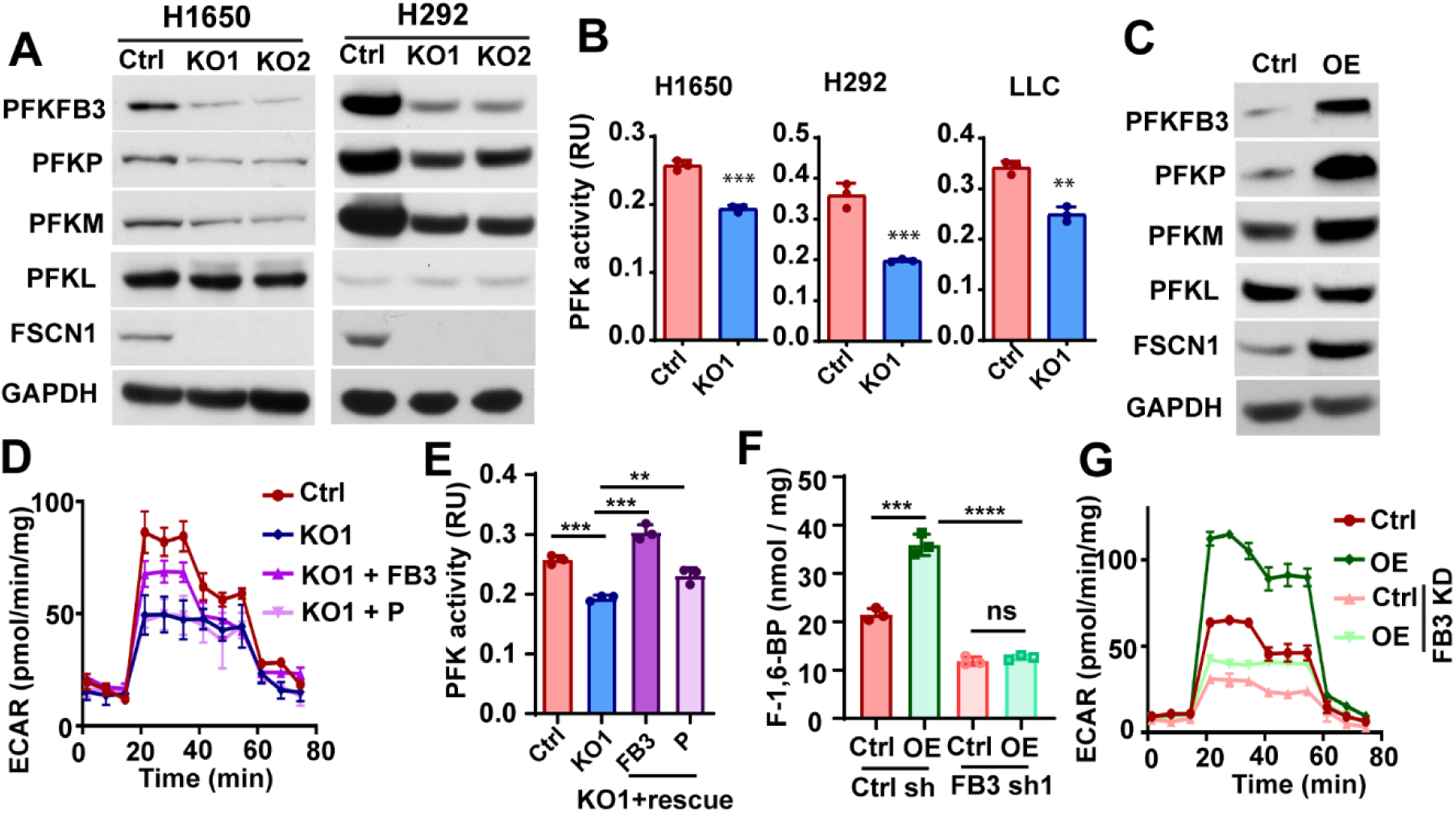
PFKFB3 is required for fascin to promote glycolysis. A and B, the effects of fascin KO on PFK protein expression levels (A) and PFK activities in NSCLC cells. C, the effect of fascin overexpression (OE) on PFK protein levels in H1650 cells. D, PFKFB3 or PFKP were ectopically expressed in fascin KO H1650 cells and the effect of ectopic PFKs on ECAR was determined. The ectopic expression of PFKFB3, but not PFKP could partially rescue ECAR in fascin KO cells. E and F, the effect of ectopic PFKFB3 (FB3) or PFKP (P) on PFK activities (E) and F-1,6-BP levels in fascin KO H1650 cells. G, the effects of PFKFB3 knockdown (FB3 KD) on ECAR in H1650 cells stably expressing control vector (Ctrl) or wild type fascin (OE). Data in B, E and F were analyzed by two-tailed, two-sample unpaired Student’s test. Representative results from at least three independent experiments are shown. **, *** and **** indicated p<0.01, 0.001 and 0.0001, respectively. ns, not significant.

Intriguingly, the overexpression of fimbrin, another actin bundling protein, had no effect on PFK protein upregulation (Fig. S2A). To determine the role of PFK2 and PFK1 in fascin-mediated glycolysis, we ectopically expressed PFKFB3 and PFKP in fascin KO cells (Fig. S2B). As shown in Fig. 3D and S2C-S2D, the ectopic PFKFB3 was able to partially restored glycolysis and lactate production in fascin KO cells. The ectopically expressed PFKP had no effect on ECAR or lactate in fascin KO cells (Fig. 3D and S2C-S2D). PFKFB3 ectopic expression was able to restore PFK activities in fascin KO cells to a level similar to control cells (Fig. 3E). In contrast, the effect of ectopic PFKP on PFK activities was marginal (Fig. 3E). To further determine whether PFKFB3 might be required for fascin-mediated glycolysis, we used shRNA to knockdown PFKFB3 in H1650 cells (Fig. S2E). As shown in Fig. 3G and Fig.S2F, PFKFB3 KD abrogate the ability of fascin to increase FBP production and greatly diminished the abilities of fascin to increase ECAR and lactate. Taken together, our data suggested that fascin promote glycolysis in NSCLC by upregulating PFKs and PFKFB3 is critical for fascin to promote glycolytic metabolism.

### Fascin activates the transcription of PFKFB3 through a YAP1/TEAD binding site on its promoter

The protein stability of PFK1 and PFK2 could be regulated through proteasomal degradation (26). However, we found that treatment of control and fascin KO cells with proteasome inhibitor MG132 had no effect on PFKFB3 levels in H1650 cells (Fig. S3A). Instead we noted upregulation of PFKFB3 mRNAs in fascin OE cells and downregulation of PFKFB3 transcripts in KO cells (Fig. S3B). Therefore, we entertained the hypothesis that fascin might promote glycolysis through transcriptional activation of PFKFB3.

Recently the Hippo pathway transcription co-activator YAP1 emerged as an important sensor of actin cytoskeleton tension and regulator of glycolysis in cancer (27). The actin cytoskeleton remodeling could activate YAP1 and its paralog TAZ through mechanisms that are still not completely understood (28). We observed that fascin KO decreased the protein levels of YAP1 (Fig. 4A and Fig. S3C) while increasing the levels of pS127-YAP1 (Fig. 4A). Fascin KO also increased the levels of active LATS1, the serine/threonine kinase responsible for phosphorylation of YAP1 at S127 and promoting its degradation (Fig. 4A), while suppressing the nuclear localization of YAP1 (Fig. 4B-4C). The expression levels of TAZ was not affected by fascin KO (Fig. S3C). Fascin KD decreased the expression of YAP1 and PFKFB3, which could be restored by WT but not by bundling-defective mutants of fascin (Fig. 4D). The protein levels of TAZ was not affected by fascin KD or rescue (Fig. 4D). Taken together, our data suggested that fascin overexpression in NSCLC activates YAP1 and increases PFKFB3 expression through its actin crosslinking activity, likely through LATS1.

**Figure 4.**
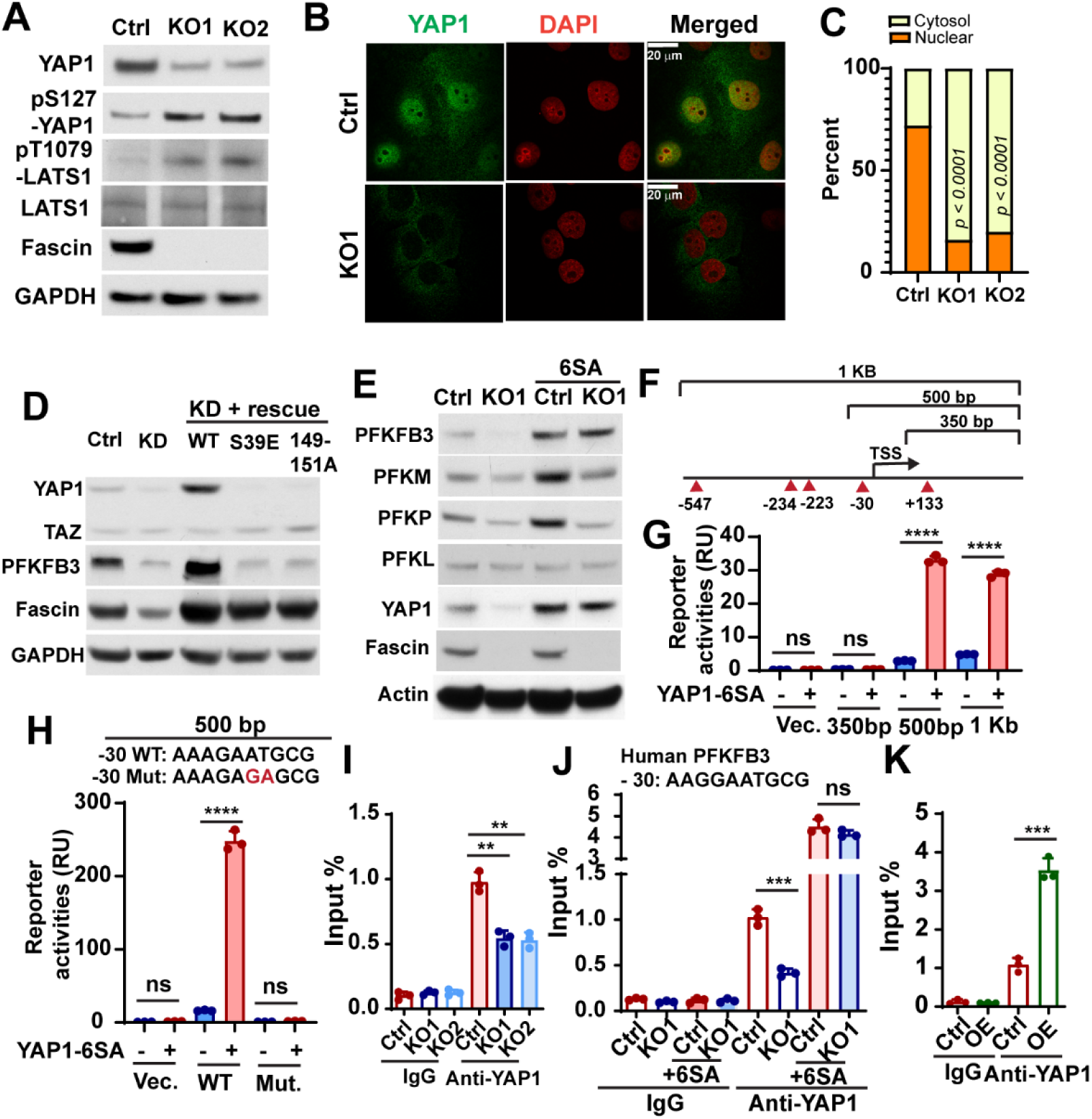
Fascin promote the transcription of PFKFB3 by activating YAP1. A, Western blotting showing the effect of fascin KO on the levels of total YAP1 protein, phosphorylated YAP1 (pS127), phosphorylated LATS1 (pT1079) and total LATS1 in H1650 cells. B and C, representative images (B) and quantitation (C) showing YAP1 nuclear translocation were inhibited in H1650 fascin KO cells. n=50 cell per group. D, the effect of fascin knockdown (KD) or the rescue of the knockdown (with ectopic wild-type fascin (WT) or the bundling defective mutants (S39E and 149-151A)) on YAP1 and PFKFB3 protein levels in H1650 cells. E, the effect of constitutively active YAP1 mutant (6SA) on the protein expression levels of PFKFB3, PFKM, PFKP and PFKL in control or fascin KO1 H1650 cells. F, schematic illustration of *Pfkfb3* promoter reporter constructs and the position of TEAD binding motifs in the promoter. G and H, the effect of YAP1-6SA mutant on the transcription of luciferase when cotransfected with indicated luciferase reporters, as determined by the dual-luciferase reporter assay. The ability of YAP1-6SA to promote luciferase transcription was abolished by the GA mutations in the 500 bp reporter construct (H). I- J, Chromatin-immunoprecipitation (ChIP) assay indicates fascin KO could decrease the binding of YAP1 to murine (I) or human (J) *PFKFB3 / Pfkfb3* promoter in LLC or H1650 cells. The binding of YAP1-6SA mutant to PFKFB3 promoter was not affected by fascin KO (J). K, ChIP assay showing that fascin overexpression could increase the YAP1 binding with PFKFB3 promoter in H1650 cells. Data in G-K were analyzed by two-tailed, two-sample unpaired Student’s test. Data in C were analyzed by two-tailed Fisher’s exact test. Representative results from at least three independent experiments are shown. **, *** and **** indicated p<0.01, 0.001 and 0.0001, respectively. ns, not significant.

To determine whether fascin might regulate PFKs through YAP1, we expressed YAP1-6SA, a constitutively active mutant of YAP1, in control or fascin KO cells. As shown in Fig. 4E, YAP1- 6SA was able to increase the protein levels of PFKFB3, PFKM and PFKP in control cells and restore the expression of PFKFB3 in KO cells to levels comparable to control-6SA cells (Fig. 4E). However the protein expression levels of PFKM and PFKP in KO1-6SA cells were still much lower than in control-6SA cells (Fig. 4E). These data suggested that fascin upregulate PFKFB3 expression through YAP1, while the upregulation of PFKM and PFKP by fascin likely also involved a YAP1-independent mechanism.

YAP1 regulate the transcription of its target genes by forming complex with TEAD transcription factors (27). Our survey of the JASPAR database (29) indicated that there are multiple TEAD binding motifs in the proximal promoter region of both murine and human PFKFB3 gene. To determine whether YAP1 might directly activate the transcription of PFKFB3, we constructed several luciferase reporters including different promoter regions from the *Pfkfb3* gene (Fig. 4F). As shown in Fig. 4G, the ectopic expression of YAP1-6SA was able to robustly activate the transcription of both 1KB and 500bp reporter but not the 350bp construct (Fig. 4G). This led us to further examine a single TEAD1/4 binding motif located 30bp upstream of the transcription start site (TSS). Mutation of this motif from AAAGAATGCG to AAAGAGAGCG completely abrogated the ability of YAP1-6SA to activate luciferase transcription in the 500bp reporter (Fig. 4H). The −30bp TEAD1/4 binding motif is conserved between human and murine (AAGGAATGCG in human). To determine whether fascin might promote the binding of YAP1 to this TEAD1/4 binding motif, we used chromatin-immunoprecipitation (ChIP) assay to examine the binding of YAP1 to this site in control and KO cells. As shown in Fig. 4I-4J, fascin KO decreased the binding of YAP1 to this site in the promoter of both murine and human PFKFB3 genes. In contrast, the binding of ectopically expressed YAP1-6SA mutant was not affected by fascin KO (Fig. 4J). Conversely, there was more than 3-fold increase in the binding of YAP1 to *PFKFB3* promoter when fascin was overexpressed in H1650 cells (Fig. 4K). Taken together, our data support that *PFKFB3* is a direct YAP1/TEAD target gene and fascin activates the transcription of *PFKFB3* through a YAP1/TEAD binding site on its promoter.

### Fascin upregulation correlates with YAP1 activation and PFKFB3 overexpression in cancer patients across different types of cancer

We have previously examined the expression of fascin and YAP1 through immunohistochemistry staining in the same cohort of 113 lung adenocarcinoma patients in two separate studies (7, 30). A review of these staining results indicated that lung cancer patients with high expression levels of fascin also tend to have strong nuclear staining of YAP1 (*r=0.497, p<0.001, n=113,* Spearman’s rank correlation test) (Fig. 5A-5B), which implicated the activation of YAP1 by fascin in lung cancer patients. To further evaluate the notion that fascin might regulate the YAP1-PFKFB3 circuit to promote glycolysis in cancer patients, we examined the correlation between multiple PFK1, PFK2 genes and two well-established YAP1 target genes (CTGF and CYR61) in the TCGA RNA sequencing database (Fig. 5C-5D, Fig. S4 and Table S1). Fascin mRNA levels significantly correlated with PFKFB3 and PFKP in lung cancer (LUAD, lung adenocarcinoma and LUSC, lung squamous cell carcinoma) and many other types of cancer, most notably in liver hepatocellular carcinoma (LIHC), colon adenocarcinoma (COAD) and rectum adenocarcinoma (READ) (Fig. 5C, Fig. S4A-S4B and Table S1). Significant correlation between fascin and YAP1 target genes were also observed in LUAD, READ, LIHC and COAD (but not in LUSC), among other types of cancer (Fig. 5D, Fig. S4C- S4D and Table S1). Taken together, our data indicated that fascin might be able to promote glycolysis and activate YAP1 signaling through a conserved mechanism across different types of cancer.

**Figure 5.**
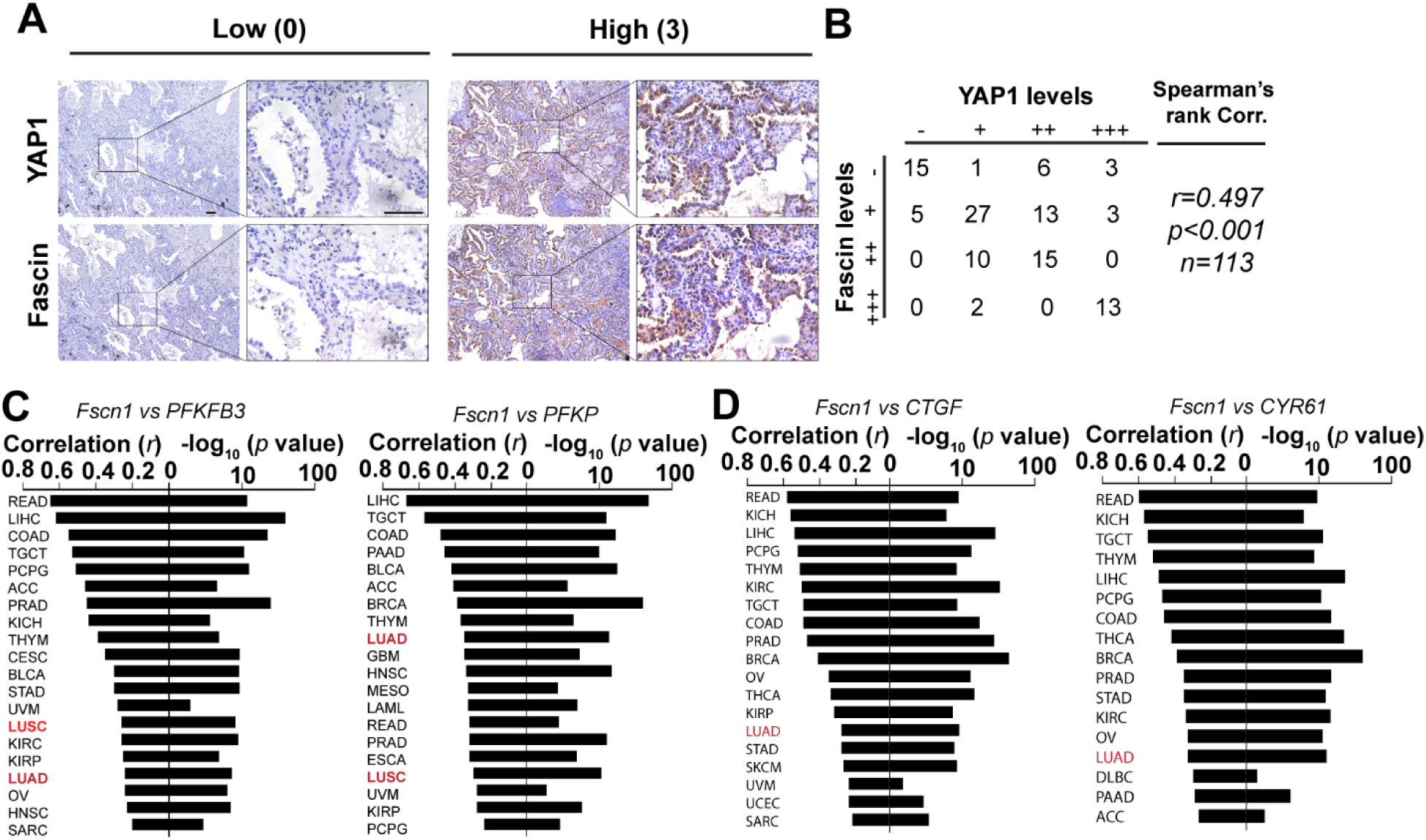
Fascin expression correlates with PFKs and YAP1 target genes in cancer patients. A, representative immunohistochemistry staining images showing expression levels of YAP1 and fascin in lung adenocarcinoma patients. B, correlation between fascin and YAP1 staining intensities in a cohort of 113 lung adenocarcinoma patient. C, waterfall plot showing the correlation between fascin and PFKFB3 or PFKP in different cancers from the TCGA RNA sequencing database. D, waterfall plot showing the correlation of fascin and YAP1 target genes (CTGF and CYR61) in different cancers from the TCGA RNA sequencing database. Correlation coefficient (*r*) and *p* value in B-D were determined using Spearman’s rank test.

### PFKFB3 is required for fascin to promote lung cancer growth and metastasis

To understand glycolytic functions of fascin in lung cancer metastasis, we knockdown PFKFB3 in control or fascin OE H1650 cells. Luciferase-labeled Control or fascin OE cells expressing control shRNA or PFKFB3 shRNA were orthotopically injected into the left lung of nude mice. The growth of tumors was monitored by bioluminescence imaging (BLI) every week (Fig. 6A and 6B). 30 days after implantation, mice were euthanized and the development of metastasis was evaluated through *ex vivo* BLI imaging (Fig. 6C-6H). Fascin OE in control shRNA H1650 cells increased the BLI signal by more than 2-fold (Fig. 6A and 6B). The knockdown of PFKFB3 inhibited BLI signal by more than 95% and fascin OE was not able to increase BLI signal in PFKFB3 shRNA group (Fig. 6A-6B). *ex vivo* BLI imaging further confirmed that PFKFB3 was required for fascin to promote H1650 tumor growth and metastasis in this model (Fig. 6C-6H). Fascin overexpression increased the BLI signals of primary tumor (left lung) (Fig. 6C-6D) and metastasis to the contralateral lung (right lung) (Fig. 6E-6F) by approximately 3 and 6 fold, respectively. The knockdown of PFKFB3 decreased *ex vivo* BLI signals in both lungs by more than 99% when compared to the control shRNA group; the ectopic expression of fascin was not able to increase BLI signals in either primary tumor or contralateral lung metastases in H1650 tumor expressing PFKFB3 shRNA (Fig. 6C-6F). We also observed chest wall metastases in 50% or 100% of control shRNA H1650 group with or without fascin overexpression, respectively (Fig. 6G and 6H). No chest wall metastases was observed in any mice in the two groups expressing PFKFB3 shRNA (Fig. 6H). Many of these chest wall metastases, especially those in fascin OE group, were visible to the naked eye (Fig. S5A, yellow arrow) and could be confirmed by H&E staining (Fig. S5B).

**Figure 6.**
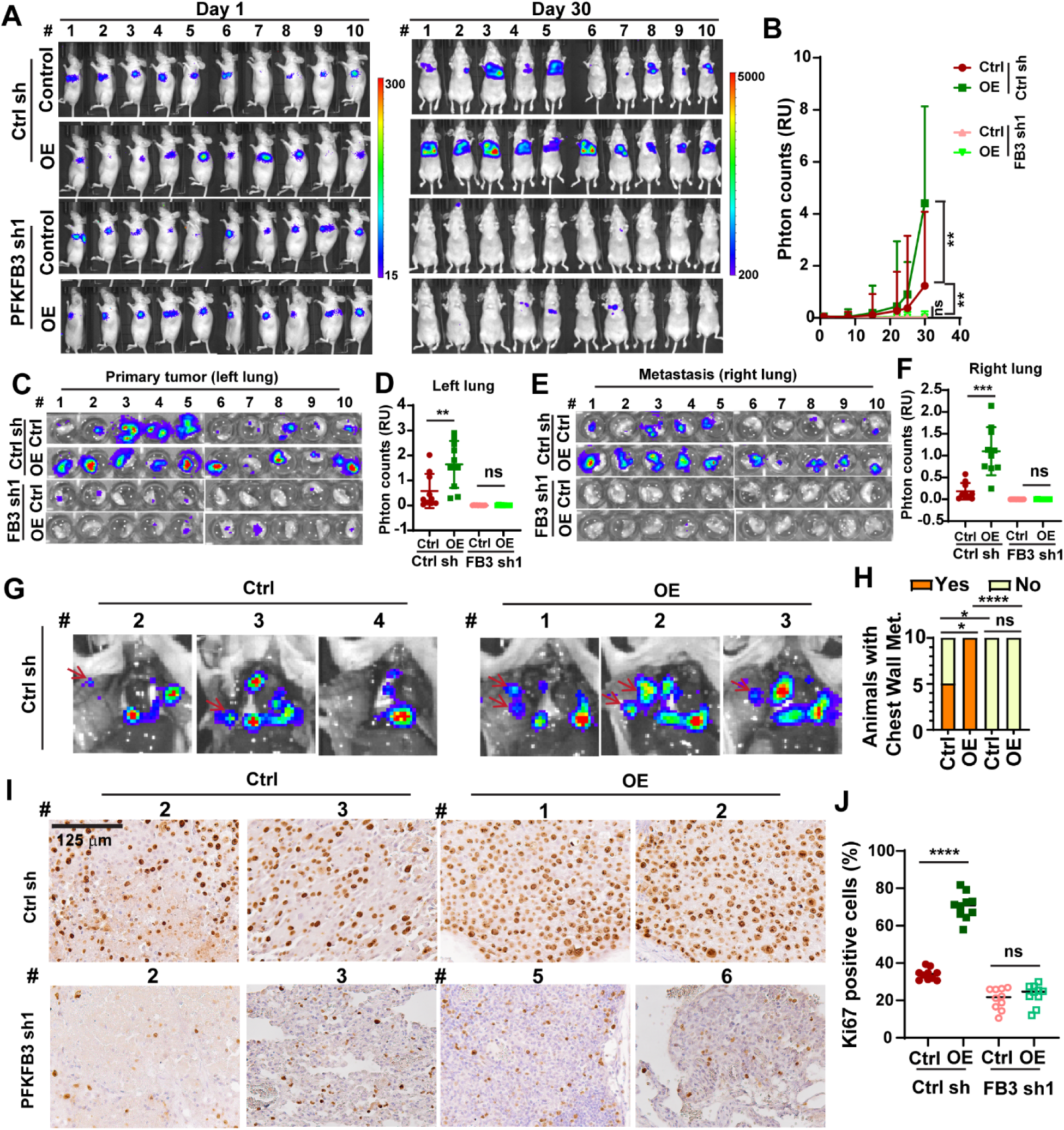
PFKFB3 is required for fascin to promote tumor growth and metastasis in lung cancer. A and B, the effects of fascin overexpression (OE) and / or PFKFB3 knockdown on orthotopic lung cancer growth and metastasis. A, BLI images of all the mice on day 1 and day 30 after orthotopic injection. B, quantitation of BLI imaging data from orthotopic experiment in A. n=10 female nude mice per group. C-F, *ex vivo* bioluminescence imaging of extracted left lungs (C and D) or right lungs (E and F). D and F are quantitation of imaging data in C and E, respectively. G, representative *ex vivo* bioluminescence imaging of the opened chest (G) showing metastases lesion on the chest wall (red arrow) in control shRNA group mice with or without fascin overexpression. H, summary of the number of mice in each group with chest wall metastases. n = 10 mice per group. I and J, representative Ki67 immunohistochemistry staining images (I) and quantitation of Ki67 positive cells (J) showing that fascin overexpression increased the proportion of Ki67 positive cells, which was abrogated by the shRNA knockdown of PFKFB3. n = 10 mice per group. Data in B were analyzed using two-tailed two-way ANOVA. D, F, and J were analyzed by two-tailed, two-sample Mann Whitney test. Data in H was analyzed using Chi-square test. *, **, *** and **** indicated p<0.05, 0.01, 0.001 and 0.0001, respectively. ns, not significant.

We and others have previously reported that NSCLC patients with elevated levels of fascin in NSCLC also had larger tumor size and higher percentage of Ki67 staining, a marker for proliferating cells (7, 31). Therefore, we evaluated the effects of fascin overexpression and PFKFB3 knockdown on Ki67 staining in orthotopic xenograft tumors. As shown in Fig. 6I and 6J, overexpression of fascin increased Ki67 positive cells from 34% to 70%. The knockdown of PFKFB3 decreased the Ki67 positive cells to 21%, and fascin OE in PFKFB3 shRNA tumor was not able to increase the proportion of proliferating cells in orthotopic tumor. Taken together, our data supported that PFKFB3 is required for fascin to promote lung cancer growth and metastasis.

### Pharmacological inhibition of fascin could be used to suppress YAP1-PFKFB3 signaling and glycolysis in NSCLC

Since fascin-mediated upregulation of YAP1 and PFKFB3 depends on its actin bundling activity, we reasoned that pharmacological fascin inhibitors such as G2 could be used to suppress glycolysis in lung cancer. As shown in Fig. 7A and 7B, G2 remarkably reduced the levels of PFKFB3 and YAP1 in lung cancer cells starting from 20-40 µM. G2 treatment had no effect on the protein levels of either PFKFB3 or YAP1 in fascin KO H1650 cells, which indicated that the G2 effect was due to its inhibition of fascin (Fig. S6A). Since fascin expression strongly correlates with PFKFB3 and YAP1 target genes in liver cancer and colorectal cancer in the TCGA database, we also examined the effects of G2 on these cancer cells. As shown in Fig. 7C, G2 treatment robustly suppressed the expression of YAP1 and PFKFB3 in liver and colorectal cancer cells as well. G2 treatment also inhibited both ECAR and OCR in H1650 (Fig. 7D-7G), which recapitulated the effects of fascin KO or KD in these cells.

**Figure 7.**
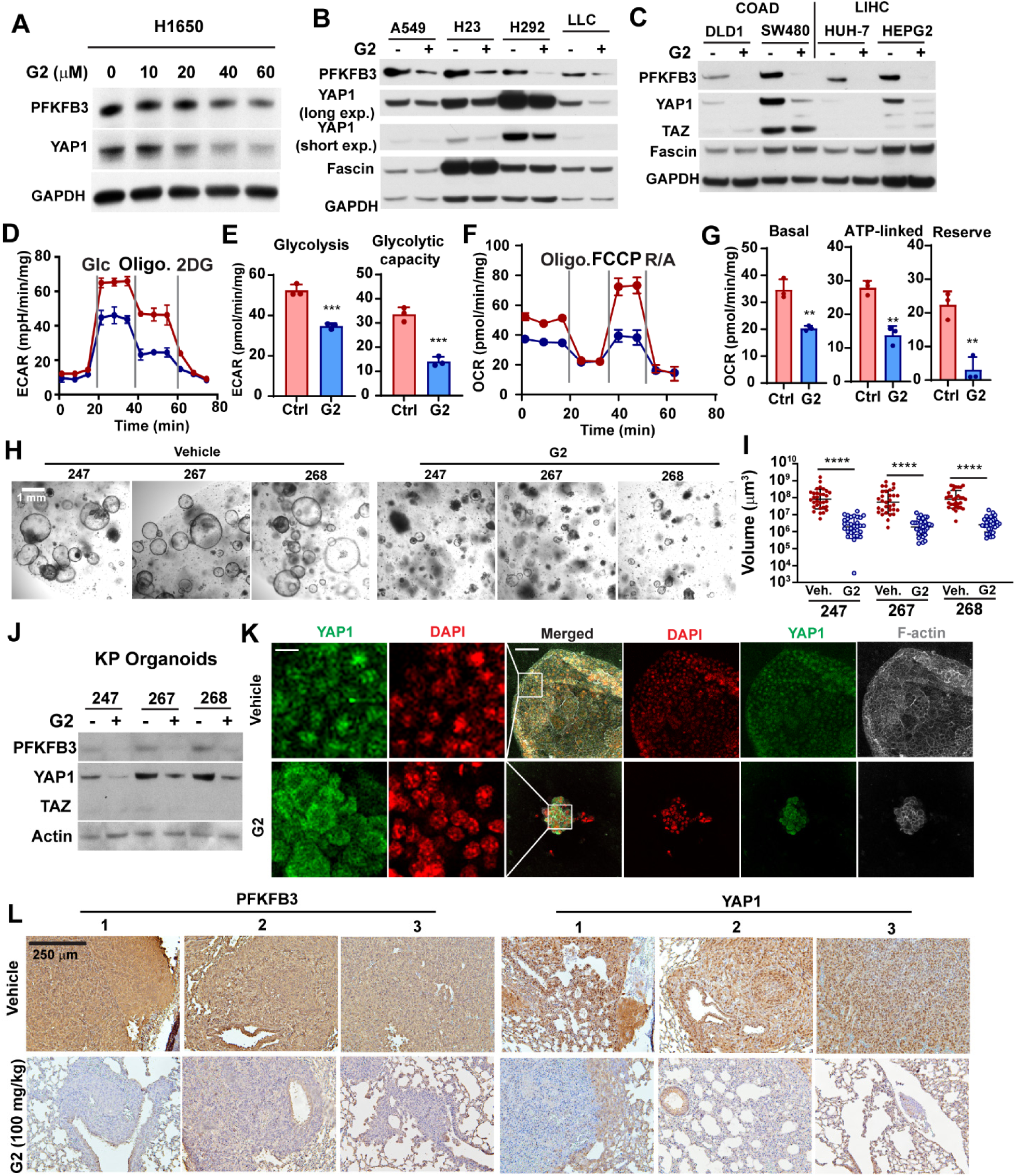
Pharmacological inhibition of fascin could be employed to suppress YAP1 and PFKFB3 in lung cancer. A-C, Western blotting showing the effect of G2 treatment on the protein levels of PFKFB3 and YAP1 protein in NSCLC cells (A and B) or liver and colorectal cancer cells (C). D-G, the effect of G2 treatment on ECAR (D and E) and OCR (F and G) in H1650 cells. E and G are quantitation of ECAR and OCR measurement in D and F, respectively. H and I, representative bright field images of tumoroids (H) and quantitation of tumoroid size (I) showing the effects of G2 treatment on tumoroid growth. Tumoroids were derived from lung tumors extracted from 3 KP mice and treated with vehicle or 40 µM G2 for 48 hours before imaging. 30 organoids were measured for each group. J, western blotting showing the effect of G2 treatment (40 µM, 48 h) treatment with on PFKFB3 and YAP1 protein levels in KP organoids. K, representative immunofluorescence images showing YAP1 subcellular localization in tumoroids treated with G2 or vehicle control. L, representative immunohistochemistry staining images showing expression levels of PFKFB3 and YAP1 in LLC tumors in mice treated with vehicle or G2 (100mg/kg daily *via* i.p.). Data in E and G were analyzed by two-tailed, two-sample unpaired Student’s test. Data in I were analyzed by Mann Whitney test. Representative results from at least three independent experiments are shown (A-K). *, **, *** and **** indicated p<0.05, 0.01, 0.001 and 0.0001, respectively.

Patient and GEMM (genetically engineered mouse model)-derived organoid cultures are able to recapitulate the characteristics of tumors they were derived from and could be used to predict the response to cancer therapies (32). *Kras*^LSL-G12D^; *p53^fl/fl^* (KP) model was able to faithfully recapitulate characteristics of lung adenocarcinoma in human (33). Therefore, we tested the effects of G2 treatment on the growth of tumoroid culture prepared from the KP lung cancer model. The formation of lung cancer in KP mice were induced by intratracheal instillation of adenoviral Cre-recombinase (33). Three months after viral induction, lung cancer tissues were harvested for the preparation of tumoroids. G2 treatment dramatically decreased tumoroid volume by 40 to 50-fold (Fig. 7H and 7I) and suppressed the protein expression levels of PFKFB3 and YAP1 in tumoroid culture from all three mice (Fig. 7J). Immunofluorescence staining revealed that G2 treatment induced the translocation of YAP1 from nuclear to the cytosol in KP tumoroids (Fig. 7K).

We have previously used G2 to treat nude mice tail-vein injected with LLC cells (7). G2 treatment significantly reduced the tumor burden in the lung of mice injected with LLC cells via tail vein (Fig. S6B) and improved the survival of these animals (Fig. S6C). IHC staining in tumor tissues harvested from these mice also showed that LLC tumors from G2 treated mice had marked decrease in both YAP1 staining and PFKFB3 staining (Fig. 7L). Intriguingly, the expression levels of YAP1 and PFKFB3 in adjacent normal lung tissue appeared to be unaffected, which indicated that G2 could be used to specifically suppress YAP1 signaling and glycolysis in cancer cells with fascin upregulation. Taken together, our data indicated that pharmacological inhibitors of fascin could be used to suppress glycolytic metabolism, tumor growth and metastasis in lung cancer and potentially in other types of cancer with high fascin expression levels.

## Discussion

Fascin is one of the most frequently upregulated actin binding protein in metastatic cancer (1, 2). The overexpression of fascin in metastatic cancer was due to inflammatory cytokines in the tumor microenvironment, epithelial-to-mesenchymal transition program and environmental stresses such hypoxia and nutrient limitation (1, 34, 35). Although it is believed that fascin promotes tumor progression by increasing tumor cell migration and invasion, fascin expression in NSCLC correlates with larger tumor size and increase in Ki67 staining (7, 31). The inducible deletion of fascin in lung cancer cells prevents the colonization of lung by disseminated cells (7). These findings suggested a role for fascin in regulating cancer cell proliferation and metastatic colonization. Our data here, together with our earlier report (7), supported a metabolic role of fascin in lung cancer and possibly other types of cancer. By remodeling mitochondrial actin filament and activatiing the YAP1-PFKFB3 signaling circuit, fascin augments both mitochondrial OXPHOS and glycolysis to increase metabolic stress resistance and cell proliferation under glucose-limited conditions. It is possible that fascin promote lung cancer growth and metastatic colonization by increasing the metabolic flexibility. Consistent with this notion, fascin overexpression increased the proportion of Ki67 positive cells in the orthotopic tumor and fascin-mediated proliferation could be abrogated by shRNA depletion of PFKFB3.

The shRNA depletion of PFKFB3 was also able to abrogate fascin-mediated tumor growth and metastasis in an orthotopic lung cancer xenograft model.

The remodeling of the actin cytoskeleton in mammalian cell is energetically costly. It is estimated that half of the ATP generated in the cells was consumed by the actin cytoskeleton remodeling (36). Intriguingly, many glycolytic enzymes are also well-known actin binding proteins (37). It is conceivable that the dysregulation of the actin cytoskeleton in metastatic cancer cells might affect bioenergetics and cancer metabolism. Consistent with this notion, insulin activates glycolysis by mobilizing the release and activation of actin filament-bound aldolase (20). Our data here demonstrated that fascin promotes glycolysis in NSCLC by transcriptional activation of PFKFB3. We further demonstrated that PFKFB3 is a direct target gene of the YAP1/TEAD complex and fascin overexpression in NSCLC promotes the binding of YAP1 to a conserved TEAD1/4 binding site on the PFKFB3 promoter. The upregulation of YAP1-PFKFB3 by fascin depends on its actin bundling activity. This is in contrast to earlier reports indicating the actin-bundling independent function of fascin in the regulation of Hippo pathway kinase MST1/2 (38, 39).

There is high interest in the development of anti-metastasis therapies targeting fascin. Several small molecule inhibitors for fascin bundling activities have been developed (10, 12, 13, 17). These inhibitors could be used to inhibit tumor growth and metastatic colonization and to reprogram cancer metabolism in lung cancer and potentially other cancer with fascin upregulation.

## Acknowledgement

We would like to thank Dr. Katherine Aird for assistance with the use of YSI 2700 Biochemical Analyzer. The research in Shengyu Yang’s lab is supported by grants from the National Institute of Heath (R01CA233844), Elsa U. Pardee Foundation and a bridge grant from the Penn State Cancer Institute.

## Materials and Methods

### Cell culture

The cancer cell lines (H1650, A549, H292, LLC, DLD1, SW480, HUH-7 and HEPG2) were obtained from ATCC and authenticated using Short Tandem Repeat profiling. All cell lines used were free of microbial (including mycoplasma) contamination. H1650, H292, DLD1 and SW480 were cultured in RPMI 1640 (HyClone, SH30027.FS) medium supplemented with 10% fetal bovine serum (FBS) (Atlanta Biological, S11150) and 1% Penicillin-Streptomycin (P/S) (Gibco, 15140163). A549, LLC, HUH-7 and HEPG2 were cultured in DMEM medium (HyClone, SH30243.FS) supplemented with 10% FBS and 1% P/S. All cells were maintained at 37 °C in a humidified 5% CO_2_ incubator.

### Animal experiments

All animal experiments were performed according to protocols approved by IACUC at the Penn State College of Medicine. 7 week-old female nude mice were purchased from Charles River and housed in a pathogen-free room with a 12 h light/dark cycle. For H1650 lung colonization experiments, 5×10^6^ luciferase labeled H1650 cells were re-suspended in 60 µl RPMI medium with 5% Matrigel (serum free) and inoculated into 7 week-old female nude mice via lung orthotopic injection as previously described (30). Noninvasive bioluminescent imaging and analysis were performed as described previously using IVIS Lumina Series III (PerkinElmer).

LLC allograft tissues with G2 treatment were collected from a previous metastatic colonization study (7). 5×10^5^ luciferase labeled LLC cells stabling expressing luciferase were resuspended in in 200 µl DMEM medium (serum free) and tail-vein injected into 7 week-old Albino BL6 female mice. Mice were randomly assigned to two groups (10 mice each, female). 48 hours after tail-vein injection. Mice were administered daily with 100 mg/kg G2 (Enamine, EN300-246105) via i.p. or with vehicle control. Mice were euthanized when they reach the humane endpoint and lungs were collected for further analysis.

#### *Kras^LSL-G12D^/Trp53^fl/fl^* lung cancer model and lung organoid culture

*Kras^LSL-G12D^* (Stock No: 008179) and *Trp53^fl/fl^* (Stock No: 008462) mice were obtained from the Jackson Laboratory and crossed to generate KP mice (*Kras^LSL-G12D^/Trp53^fl/fl^*) as previously described (33). The formation of lung adenocarcinoma were induced by intratracheal instillation of Adenovirus-Cre (1×10^7^) following the protocols of DuPage et al (40). For lung intratracheal instillation, 9 weeks old KP mice was injected with 50 µl of the virus mix (10 mM CaCl_2_, 1×10^7^ VVC-U-Ad5CMVCre (from Carver College of Medicine) diluted in Opti-MEM). 3 months after viral infection, mice were euthanized and lung tumors were harvested for the preparation of organoid culture. Tumors were washed with iced cold DMEM medium 3 times and cut with a sterile scissors to approximately 1 mm^3^ pieces. The tissues were digested with collagenase Sigma, C0130) in serum free DMEM/F12 media at a final concentration of 2 mg/ml for 60 min at 37 °C. After centrifugation at 200 g for 5 minutes, the pellet was washed with medium (DMEM with 10% FBS) 5 times to rinse off collagenase. The pellet was resuspended in 300 µl of Matrigel (Corning, 354277) and plated in 24 well plate (50 µl/ well). After solidification at 37°C for 15 min, 500 µl warmed organoid culture medium per well was added (DMEF/12 500 ml, 10% FBS, 1% P/S, 5ml ITS-G (Gibico, 41400045), 10 µl A83-01 (25 mM stock, Sigma, SML0788), 100 µl Y-27632 (50mM stock, Biogems, 1293823).

### Inhibitor treatment

Unless indicated otherwise, cells or organoids were incubated with medium containing 40 µM G2 (Xcessbi, M60269) at 37 °C for 48 h in a 5% CO_2_ incubator before being used for experiments.

### Plasmids

The FSCN1 knock out was performed using pLenti CRISPR V2 vector (Addgene, 52961) encoding sgRNA targeting FSCN1 (human FSCN1 sgRNA: 1, (human FSCN1: sgRNA, GAAGAAGCAGATCTGGACGC; 2, CACCTTGAACCCGAACGCCT; mouse FSCN1: sgRNA, 1, TCGCTACCTGGCCGCCGACA; 2, AGCCGAGGCGTTCGGGTTCA). To rescue FSCN1 knockdown or overexpress FSCN1 proteins, wild type or mutant FSCN1 (S39E and 149-151A mutants) cDNAs were subcloned into pLenti.CMV.blasticidin vector (Addgene, 17486) (Campeau et al., 2009). YAP 6SA expression plasmids were obtained from Addgene (Addgene #, 42562). PFKFB3 shRNAs were from Sigma shRNA library (sh1, TRCN0000314746; sh2, TRCN0000314747) The retroviral and lentiviral particles were packaged in HEK293 cells using PEI transfection method, and concentrated as previously described (41).

### Western blotting

Western blotting was performed as the method described previously (Lin. et al., 2019). Cells were lysed in SDS-NP40 buffer (50 mM Tris, pH 8.0, 150 mM NaCl, 1% NP40, 1% SDS, 1mM protease inhibitors cocktail) on ice for 1 min. Cells were scraped from the plate and sonicated 5 sec for three times. Then lysates were then heated at 95°C for 5 min and centrifuged at 15,000 ×g, 4 °C for 10 min. To prepare cell lysate for detection of protein phosphorylation, cells were lysed in Triton X-100 buffer (50 mM Tris, pH 7.4, 150 mM NaCl, 1% Triton X-100, 1 mM EDTA, 1mM phosphatase inhibitor cocktail (Thermo Scientific, 88667), 1 mM protease inhibitors cocktail (Roche, 04693159001)) on ice for 15 min. 50 µg proteins were separated by SDS-PAGE and transferred onto polyvinylidene difluoride membrane (PVDF, Millipore, IPVH00010). The membranes were incubated in blocking buffer (5% w/v nonfat dry milk in Tris-buffered saline, 0.05% Tween-20 (TBS-T)) for 30 min at room temperature (RT), and incubated with primary antibodies over night at 4 °C, followed by incubation with secondary antibodies for 60 min at RT.

### Immunofluorescence staining and microscopy

Immunofluorescence staining was carried out as previously described (7, 42). 40,000 cells were seeded onto laminin coated coverslip in 24-well plate in for overnight. Cells were then fixed with 4% fresh paraformaldehyde (Sigma, 158127) in PBS at RT for 20 min. Cells were incubated with primary antibodies diluted in antibody dilution buffer (2% BSA 0.1% Triton X-100 in TBS) for overnight at 4 °C. Cells were then incubated with secondary antibodies for 60 min at RT, and then with DAPI (Thermo Fisher, D1306) at RT for 10 min. Extensive 5 min washing with TBS was performed between each step. After a final wash with TBS the coverslips were mounted with Fluoromount (Sigma, F4680).

For lung organoid staining, the assay was carried out as previously described with a few modifications (30). Matrigel (Corning, 354277) was thawed at 4°C overnight and used for the organoid culture. 1 × 10^4^ cells were re-suspended in 100 μl Matrigel and plated onto the glass bottom plate. After incubating at 37°C for 20 min to allow Matrigel to solidify, 2ml organoid culture medium was added. After 3 days, organoids were incubated with vehicle control of G2 (40 µM) for 48 hours. Organoids were fixed with 4% fresh paraformaldehyde in PBS at room temperature for 20 min and then incubated with YAP1 antibody in antibody dilution buffer (2% BSA 0.1% Triton X-100 in PBS) at 4°C overnight. After sequential incubation with secondary antibody (ThermoFisher, A11008) and phalloidin (ThermoFisher, A12381) for 3 hours and then DAPI for 30 min the coverslips were mounted with Fluoromount. Extensive washing with TBS was performed between each step (20 minutes each). The images were taken using Leica SP8 confocal microscope.

### Mitochondrial stress test

Oxygen consumption rate (OCR) was determined with a Seahorse XF96 Extracellular Flux Analyzer (Agilent Seahorse Bioscience) following protocols recommended by the manufacturer. 10,000 cells were seeded each well on XF96 microplates. Cells were maintained in a non-buffered assay medium (Agilent) in a non-CO_2_ incubator for 60 min before the assay. The XF Cell Mito Stress Test Kit (Agilent, 103015), was used for the assay. The baseline recordings were followed by sequential injection of following inhibitors: oligomycin (1 µM), FCCP (1 µM), and rotenone (1 µM)/ antimycin A (20 µM).

### Glycolysis stress test

Extracellular Acidification Rate (ECAR) was determined with a seahorse XF24 Extracellular Flux Analyzer (Agilent Seahorse Bioscience) following protocols recommended by the manufacturer. 50,000 cells were seeded each well on XF24 microplates. Cells were maintained in a non-buffered assay medium (Agilent) in a non-CO_2_ incubator for 60 min before the assay. The XF Cell glycolysis Test Kit (Agilent, 103020), was used for the assay. The baseline recordings were followed by sequential injection of following reagents: glucose (10 mM), oligomycin (1 µM), or 2-DG (50 mM).

### PKM2 cross-linking

3 × 10^6^ cells in a 10 cm plate were washed with 5 ml ice cold PBS 3 times. Cells were scraped from the plate in 1 ml ice cold PBS. After centrifugation at 500 × g centrifuge 5 min at 4 °C, the pellet was re-suspended in 0.5 ml ice cold PBS. Cells were frozen in lipid nitrogen and thaw at 37°C water bath for 3 times to lyse the cells. After centrifugation at 10,000 × g for 10 min at 4 °C, formaldehyde (Amresco, M134) was added to the to the supernatant to a final concentration 1%. The solution was mixed and incubated at room temperature for 20 min. 50 µg total protein was resolved in 8% non-reducing SDS-PAGE gel and blot with PKM2 antibody (Cell Signaling, 3198s).

### FBP level measurement

FBP level was determined with the kit (BioVision, K2036-100) following manufacturer’s instruction. 2 × 10^6^ cells were lysed with 500 µl ice-cold F1,6BP Assay Buffer and place on ice for 10 min. After centrifugation at 10,000 × g, 4°C for 10 min, the supernatant was transferred to a new micro-centrifuge tube and used for the assay. The fluorescence was measured at Ex/Em=535/587 nm in end point mode using FlexStation 3 (Molecular Devices).

### PFK activity assay

PFK activity was determined with a kit from BioVision (BioVision, K776) following manufacturer’s instruction. 2 × 10^6^ cells were lysed with 200 µl ice-cold Assay Buffer on ice. After centrifugation at 14,000 × g for 5 min, the supernatant was collected for protein assay. 100 µg protein in 20 µl solution were used for the assay. OD 450 nm was measured using FlexStation 3 (Molecular Devices).

### Immunohistochemistry (IHC)

IHC staining for fascin and YAP1 in patient specimens were performed in a cohort of 113 lung adenocarcinoma patients in two separate studies as previously described (7, 30). The intensity of the staining was evaluated by two independent investigators blinded to the clinic pathological data of patients using the following criteria: 0, negative; 1, low; 2, medium; 3, high. The extent of staining was scored as 0, 0% stained; 1, 1% to 25% stained; 2, 26% to 50% stained; 3, 51% to 100% stained. Five random fields (20 × magnification) were evaluated under a light microscope. The final scores were calculated by multiplying the scores of the intensity with those of the extent and dividing the samples into four grades: 0, negative (-); 1 to 2, low staining (+); 3 to 5, medium staining (++); 6 to 9, high staining (+++). The correlation between fascin and YAP1 staining intensities were determined by Spearman’s rank correlation test.

For IHC staining of YAP1, PFKFB3 and Ki67 in xenograft tissues, paraffin-embedded sections of tissues were deparaffinized and then heated in a pressure pot for 3 minutes to retrieve the antigens. The slides sections were then incubated with primary antibodies (1:200) overnight at 4°C. Antibody binding was detected using a peroxidase–conjugated secondary antibody at 37°C for 30 minutes. A DAB Substrate Kit was used to perform the chromogenic reaction.

### Chromatin immunoprecipitation (ChIP) assay

ChIP was performed with a kit from Abcam (catalog # ab500) following manufacturer’s protocols. Cells were grown to 90% confluency on a 10 cm plate. 1 × 10^6^ cells are required for each reaction. After sonication, YAP1 antibody (Cell Signaling, 14074S) was used for the immunoprecipitation overnight with rotation at 4°C. After DNA purification, the quantitative PCR was performed with the following primers:

Human *PFKFB3*

hPFKFB3_YAP1ChIP_FW: GGGATAACAGCAGCCAGGA

hPFKFB3_YAP1ChIP_RV: CTTCCACGTGGGGAGGAG

Murine *Pfkfb3*

mPfkfb3_Yap1ChIP_FW: CTGCTCTCGTCCACCCGCAG

mPfkfb3_Yap1ChIP_RV: CTCCTCTCCAGTTCTGTCGG

### Dual-luciferase report assay

Dual-luciferase report assay was determined with the kit (Promega, E1910) following manufacturer’s protocols. The promoter of murine *Pfkfb3* (350bp, 500 bp, 1000bp and 6000bp) was subcloned into pGL3 vector. Lipofectamine 2000 (ThermoFisher, 11668019) was used to transient transfect 293T cells in 12-well plate (1 µg reporter plasmid and 1 µg renilla luciferase). After 48 h, cells were lysed with 100 µl PLB buffer from the kit. 20 µl of cell lysate were transferred into 96-well reading plate and mixed with 100 µl LARII solution. Luciferase signal was read in FlexStation 3 (Molecular Devices).

### Gene Expression Profiling Interactive Analysis (GEPIA)

TCGA data analysis was performed through the GEPIA website as previously described (43).

### Metabolic flux analysis

For metabolic flux assays of glycolysis intermediates, cells were incubated with 25 mM U- ^13^C_6_ Glucose in DMEM or the control media for the indicated durations. After washing twice with PBS, 1 ml 80% methanol/water (−80℃ cooled) was added to quench and extract the metabolites.

The cells were incubated in a −80℃ freezer for at least 15 min, and then scraped into 1.7 ml tubes and centrifuged at 15000 rpm for 10 min. The supernatant was collected and dried using SpeedVac and re-dissolved in 50μL methanol/water (50/50) for LC-MS/MS analysis. LC-MS/MS analysis was performed with Waters ACQUITY UPLC systems connected to a Xevo TQ-S mass spectrometer using Selective Reaction Monitoring (SRM) methods as described previously(44, 45). Chromatographic separation was performed on a Waters UPLC BEH Amide column (2.1×150 mm, 1.7μm). Percentage contributions were calculated based on mass isotopomer measurements. FluxFix was used for the natural isotope abundance correction(46).

## Supplementary Figures

**Figure S1.**
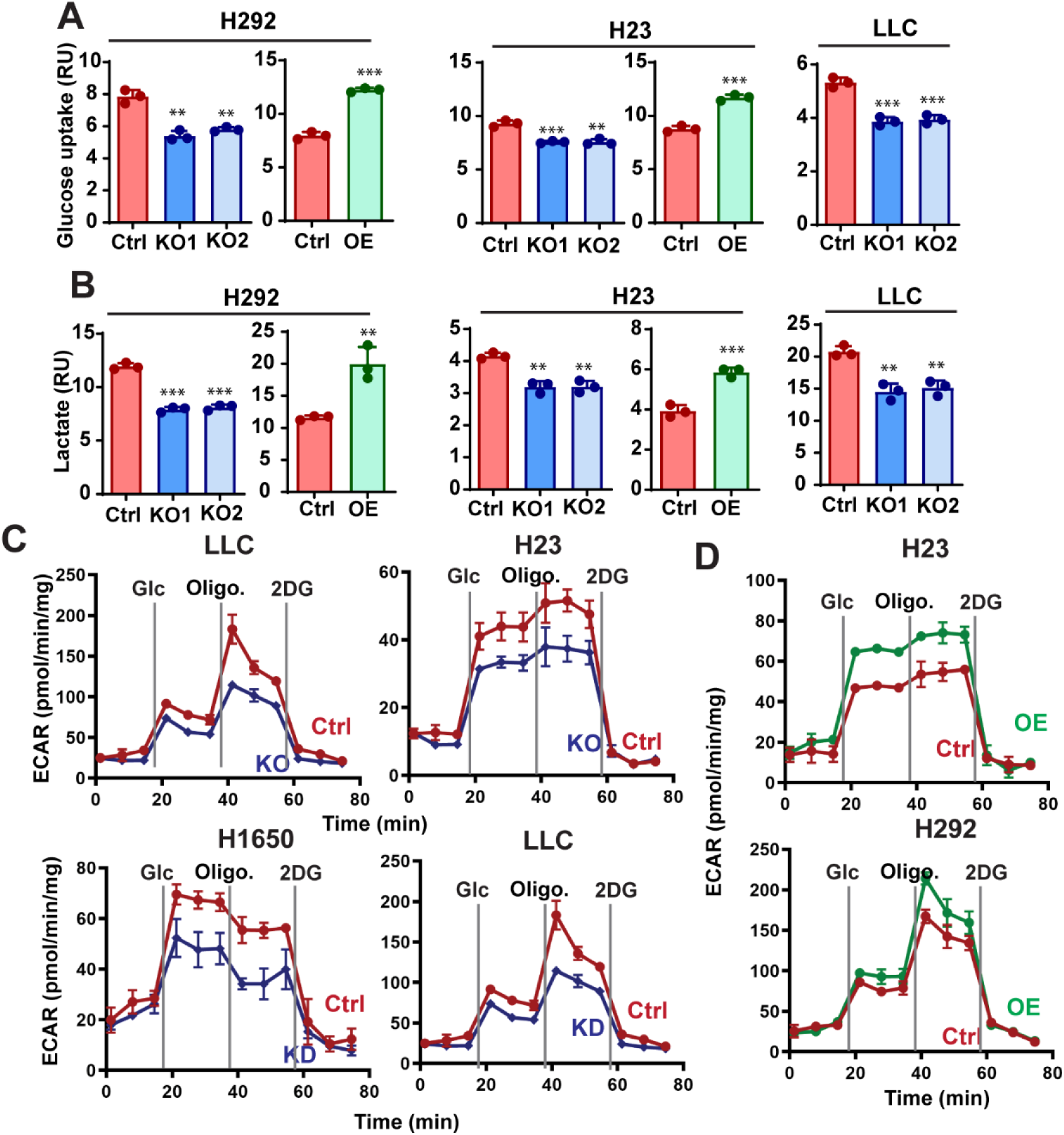
Fascin promotes glycolysis in NSCLC cells. A, the effect of fascin KO or OE on glucose uptake in NSCLC cell lines. B, the effect of fascin KO or OE on lactate production in NSCLC cell lines. C and D, the effects of fascin KO, KD (C) or OE (D) on extracellular acidification rate (ECAR), as determined by glycolysis stress test in NSCLC cell lines. Quantitation of graphs in C and D are shown in Fig. 1E and 1F, respectively. Data in A and B were analyzed by two-tailed, two-sample unpaired Student’s test. Representative results from at least three independent experiments are shown. ** and *** indicated p<0.01 and 0.001, respectively.

**Figure S2.**
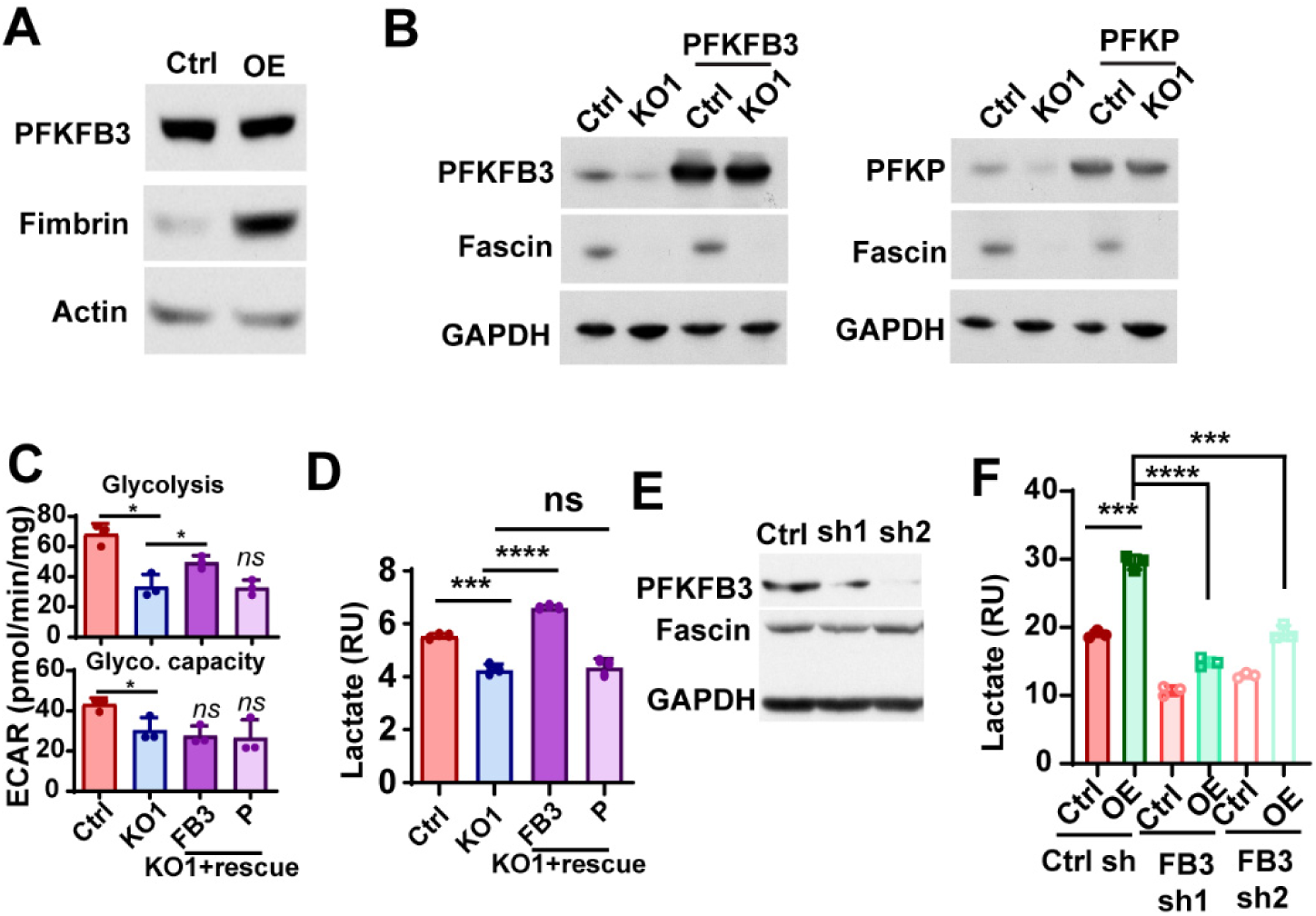
PFKFB3 regulates glycolysis. A, Western blotting showing fimbrin overexpression had no noticable affect on PFKFB3 protein level. B, Western blotting showing the expression levels PFKFB3 or PFKP in control and fascin KO1 H1650 cells with or without ectopic expression of these two proteins, respectively. C, the quantitation of ECAR in Fig 3D. The ectopic expression of PFKFB3, but not PFKP, in fascin KO H1650 cells could rescue glycolysis. D, the ectopic expression of PFKFB3, but not PFKP, in fascin KO H1650 cells was able to rescue lactate production in H1650 cells. E, Western blotting showing the effects of two independent shRNAs on the protein expression levels of PFKFB3 in H1650 cells. F, the effect of PFKFB3 shRNAs and ectopically overexpressed fascin on lactate production in H1650 cells. Data in (C, D, and F) were analyzed by two-tailed, two-sample unpaired Student’s test. These results from at least three independent experiments are shown. *, **, *** and **** indicated p<0.05, 0.01, 0.001 and 0.0001, respectively.

**Figure S3.**
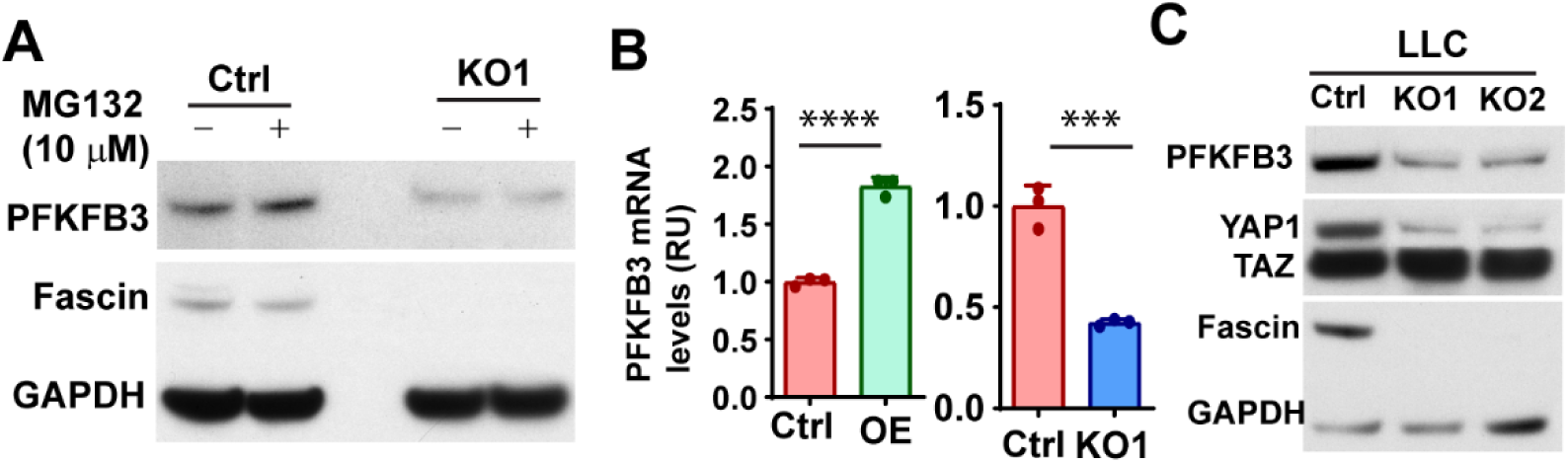
Fascin regulates PFKFB3 through transcription level. A, Western blotting showing that inhibition of proteasome with MG132 had no effect on fascin PFKFB3 protein levels in control of fascin KO H1650 cells. B, quantitative PCR results showing the expressing of PFKFB3 mRNA levels in fascin OE and fascin KO H1650 cells. C, Western blotting showing decreases in PFKFB3 and YAP1 protein levels in fascin KO LLC cells. The expression levels of TAZ was not affected by fascin KO. Data in B were analyzed by two-tailed, two-sample unpaired Student’s test. These results from at least three independent experiments are shown. *** and **** indicated p<0.001 and 0.0001, respectively.

**Figure S4.**
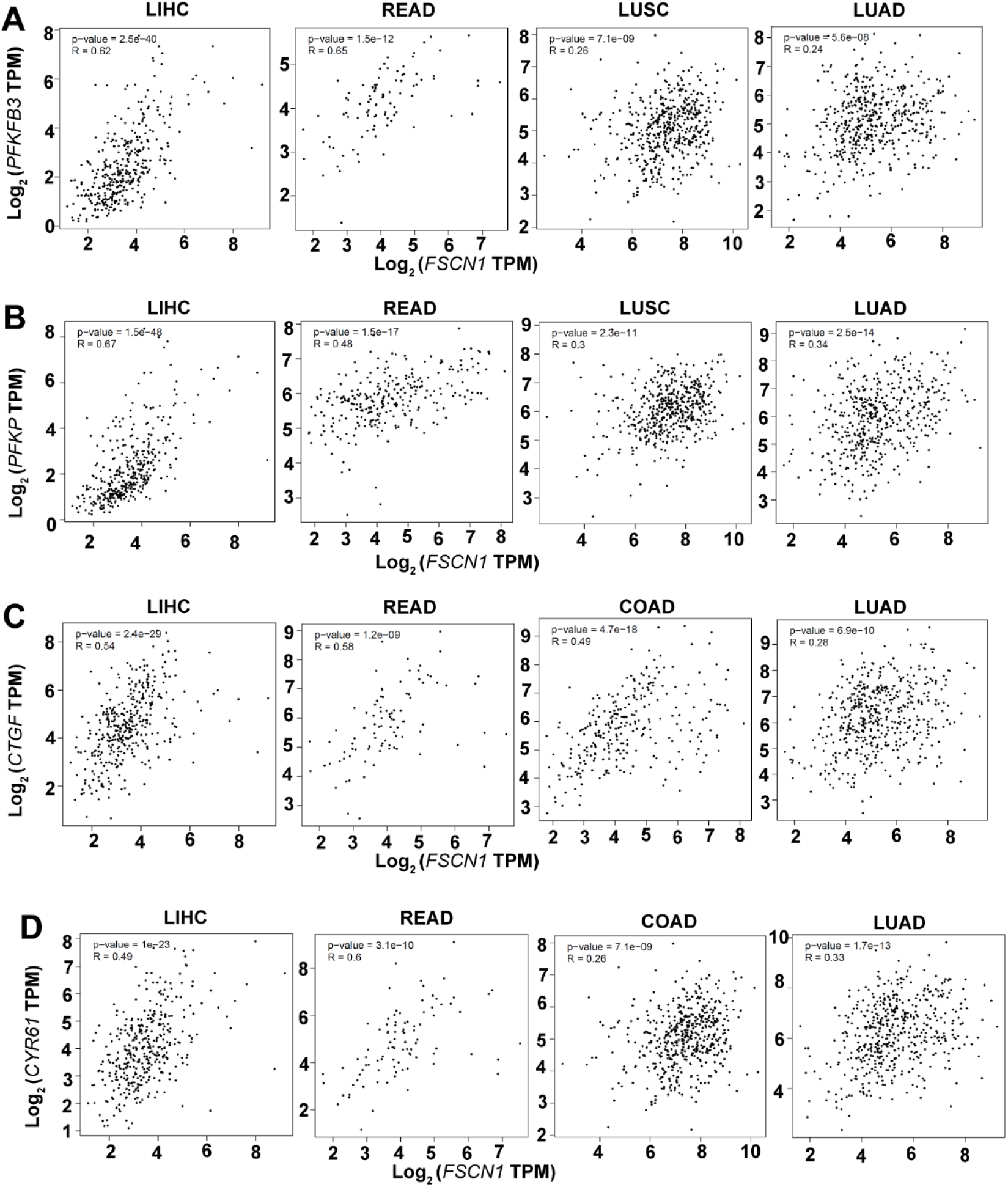
The correlation between FSCN1 mRNA expression and PFKs and YAP1 target genes in the TCGA RNA sequencing database. A and B, the correlation between FSCN1 transcript levels and PFKFB3 (A) or PFKP (B) in LIHC, READ, LUSC, and LUAD. C and D, the correlation between FSCN1 transcript levels and the mRNA levels of YAP1 target gene CTGF (C) and CYR61 in LIHC, READ, COAD, and LUAD. Data in A-B were analyzed using Spearman’s rank test.

**Figure S5.**
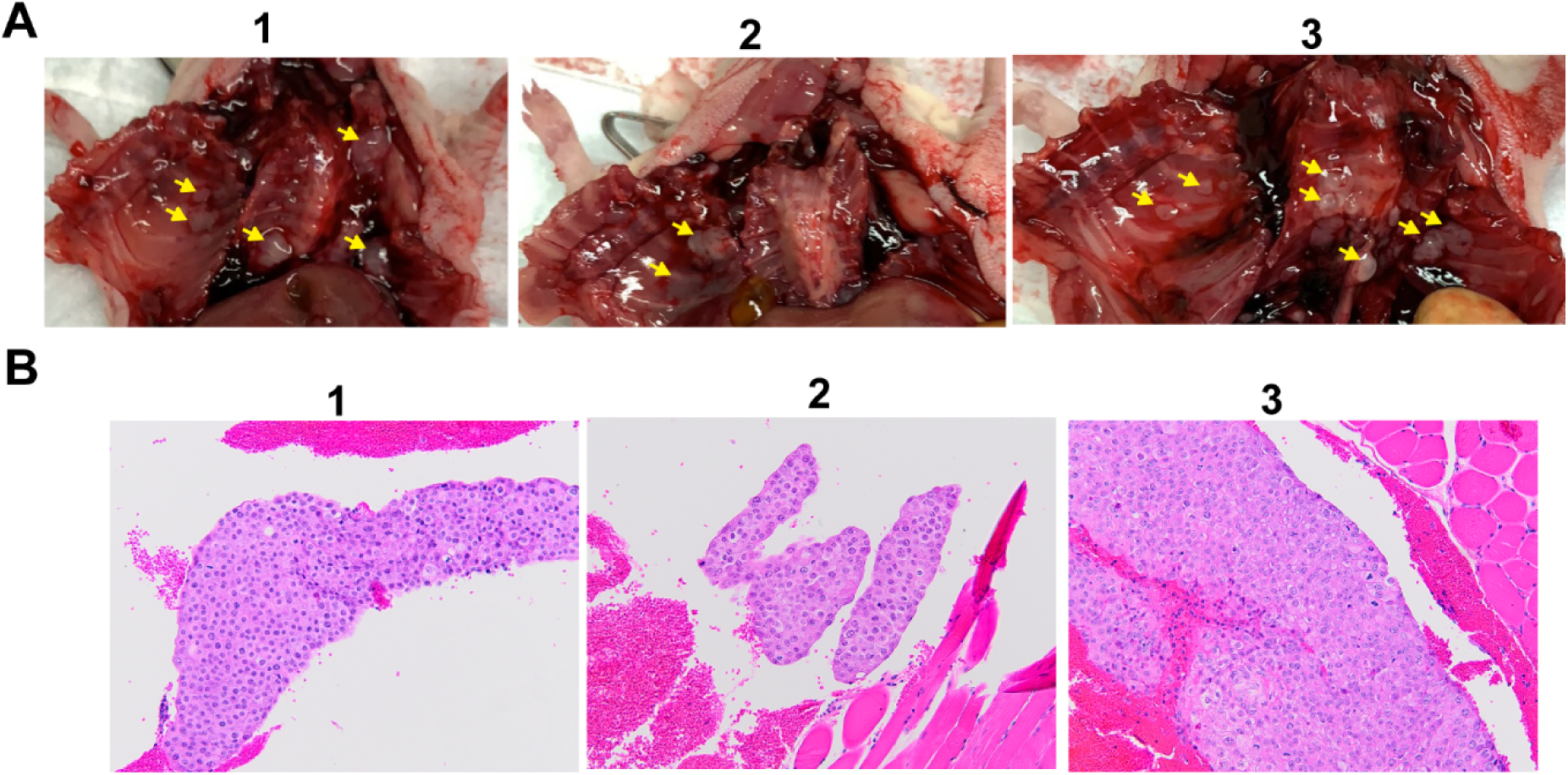
Metastatic lesions on the chest wall of mice with orthotopic H1650 xenograft. A, representative image showing the metastatic lesions (yellow arrow) on the chest wall of 3 mice orthotopically injected with H1650 cells expressing fascin and control shRNA. B, representative H & E staining images showing chest wall metastases from the 3 mice in (A).

**Figure S6.**
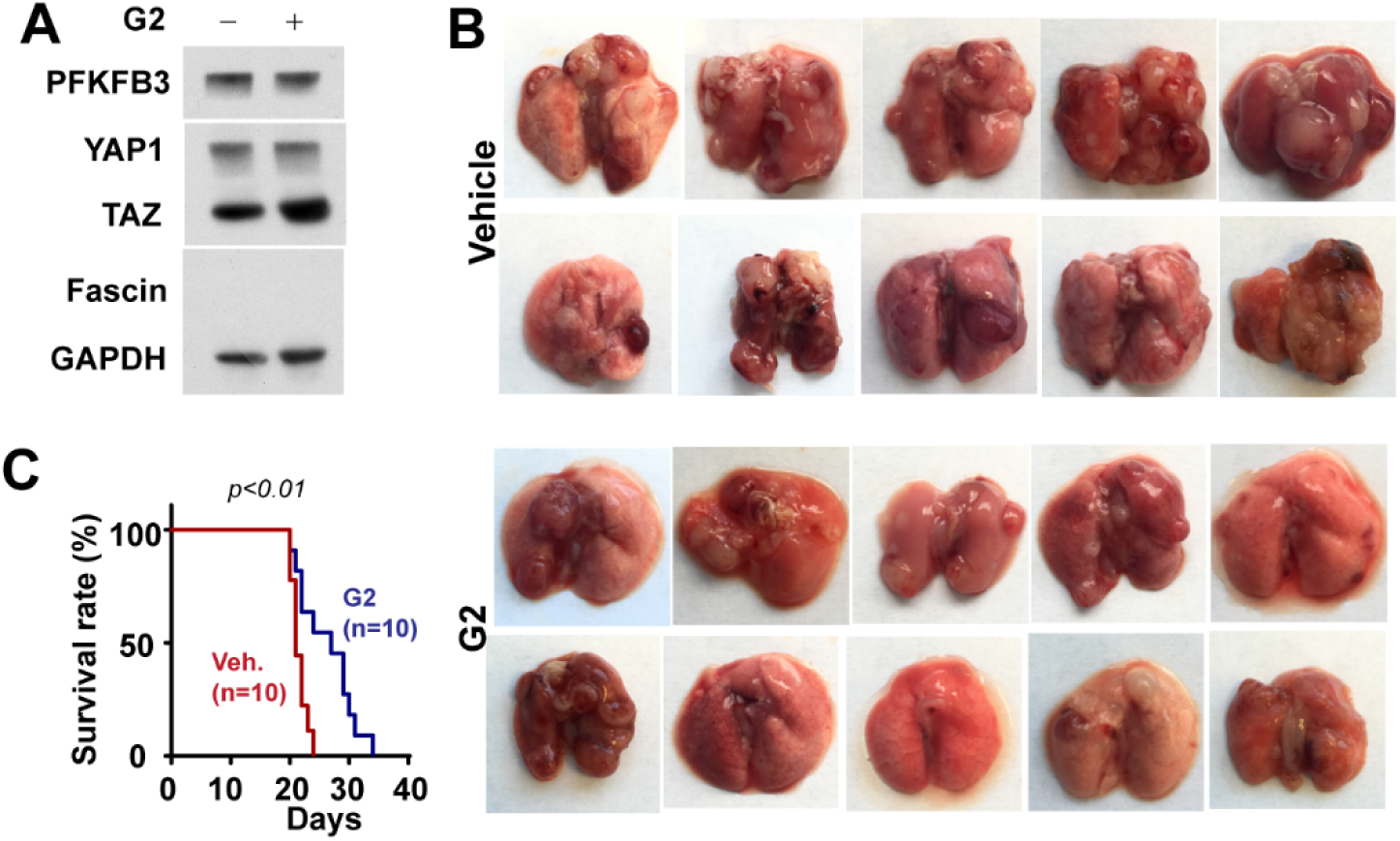
A, Western blotting showing that G2 treatment (40 µM, 48 hours) had no noticeable effect on PFKFB3 or YAP1 expression in fascin KO1 H1650 cells. B, Photos of lungs harvested from mice treated with either vehicle control or G2 (100 mg/kg, daily via *i.p.*) 2 days after injected with LLC cells via tail-vein. n = 10 mice per group. C, Kaplan-Meier survival analysis showing that G2 treatment improve the survival of animal in LLC tail-vein metastasis model.

**Table S1.**
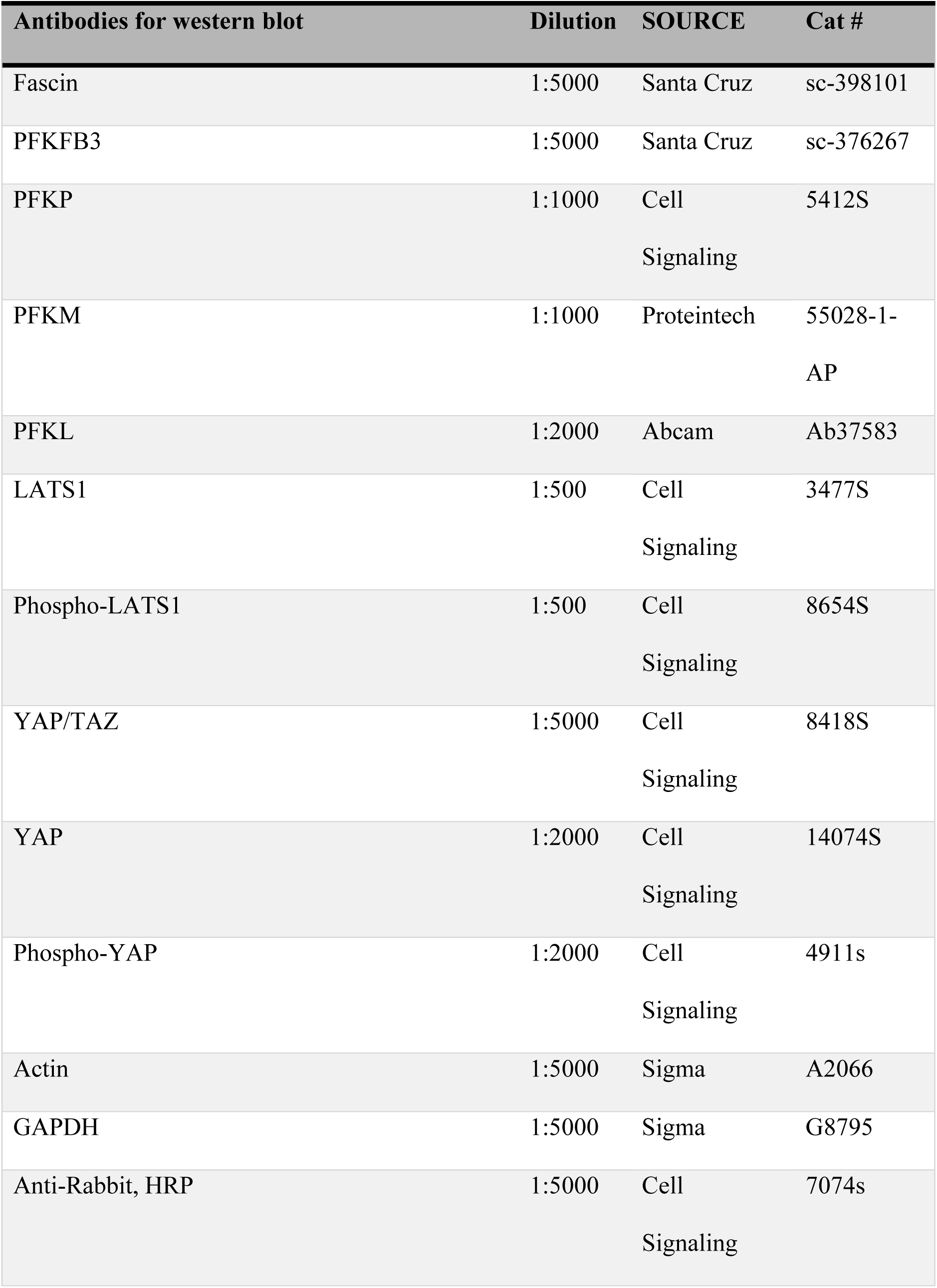
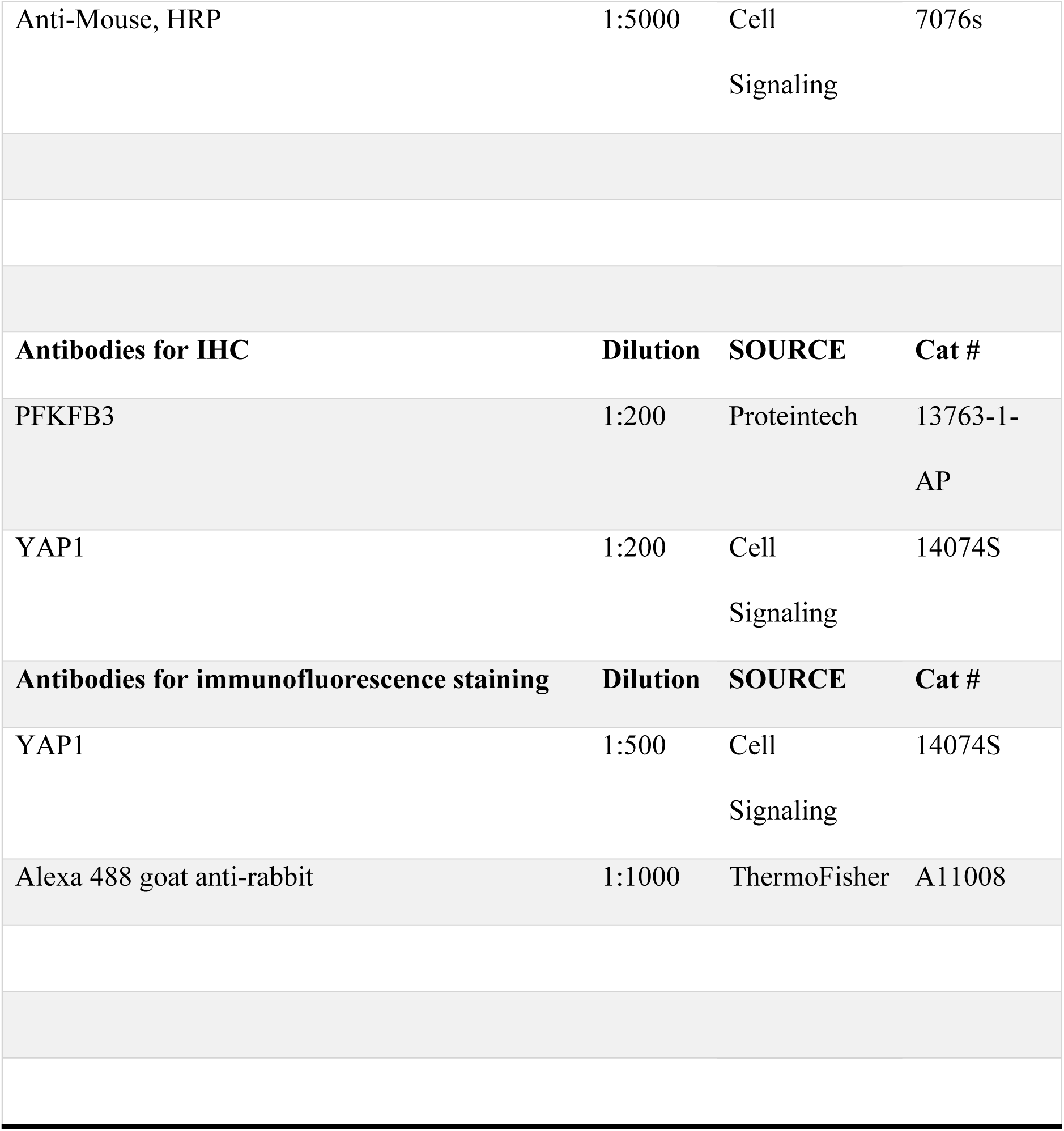
Antibodies and dilution.

## Reference

1. Lin S, Taylor MD, Singh PK, Yang S. How Does Fascin Promote Cancer Metastasis? Febs J. 2020. Epub 2020/07/14. doi: 10.1111/febs.15484. PubMed PMID: 32657526.

2. Liu H, Zhang Y, Li L, Cao J, Guo Y, Wu Y, Gao W. Fascin actin-bundling protein 1 in human cancer: promising biomarker or therapeutic target? Mol Ther Oncolytics. 2021;20:240–64. Epub 2021/02/23. doi: 10.1016/j.omto.2020.12.014. PubMed PMID: 33614909; PMCID: PMC7873579.

3. Hashimoto Y, Skacel M, Adams JC. Roles of fascin in human carcinoma motility and signaling: prospects for a novel biomarker? Int J Biochem Cell Biol. 2005;37(9):1787–804. Epub 2005/07/09. doi: S1357-2725(05)00155-X [pii] 10.1016/j.biocel.2005.05.004. PubMed PMID: 16002322.

4. Lin S, Lu S, Mulaj M, Fang B, Keeley T, Wan L, Hao J, Muschol M, Sun J, Yang S. Monoubiquitination Inhibits the Actin Bundling Activity of Fascin. J Biol Chem. 2016;291(53):27323–33. doi: 10.1074/jbc.M116.767640. PubMed PMID: 27879315; PMCID: PMC5207158.

5. Machesky LM, Li A. Fascin: Invasive filopodia promoting metastasis. Commun Integr Biol. 2010;3(3):263–70. Epub 2010/08/18. PubMed PMID: 20714410.

6. Barnawi R, Al-Khaldi S, Majed Sleiman G, Sarkar A, Al-Dhfyan A, Al-Mohanna F, Ghebeh H, Al- Alwan M. Fascin is Critical for the Maintenance of Breast Cancer Stem Cell Pool Predominantly via the Activation of the Notch Self-Renewal Pathway. Stem cells. 2016. doi: 10.1002/stem.2473. PubMed PMID: 27502039.

7. Lin S, Huang C, Gunda V, Sun J, Chellappan SP, Li Z, Izumi V, Fang B, Koomen J, Singh PK, Hao J, Yang S. Fascin Controls Metastatic Colonization and Mitochondrial Oxidative Phosphorylation by Remodeling Mitochondrial Actin Filaments. Cell reports. 2019;28(11):2824–36 e8. Epub 2019/09/12. doi: 10.1016/j.celrep.2019.08.011. PubMed PMID: 31509745.

8. Ghebeh H, Al-Khaldi S, Olabi S, Al-Dhfyan A, Al-Mohanna F, Barnawi R, Tulbah A, Al-Tweigeri T, Ajarim D, Al-Alwan M. Fascin is involved in the chemotherapeutic resistance of breast cancer cells predominantly via the PI3K/Akt pathway. Br J Cancer. 2014. Epub 2014/08/15. doi: 10.1038/bjc.2014.453. PubMed PMID: 25117814.

9. Valastyan S, Weinberg RA. Tumor metastasis: molecular insights and evolving paradigms. Cell. 2011;147(2):275–92. Epub 2011/10/18. doi: 10.1016/j.cell.2011.09.024. PubMed PMID: 22000009.

10. Han S, Huang J, Liu B, Xing B, Bordeleau F, Reinhart-King CA, Li W, Zhang JJ, Huang XY. Improving fascin inhibitors to block tumor cell migration and metastasis. Mol Oncol. 2016;10(7):966–80. doi: 10.1016/j.molonc.2016.03.006. PubMed PMID: 27071719.

11. Wang Y, Zhang JJ, Huang XY. Anti-Metastasis Fascin Inhibitors Decrease the Growth of Specific Subtypes of Cancers. Cancers (Basel). 2020;12(8). Epub 2020/08/23. doi: 10.3390/cancers12082287. PubMed PMID: 32824026.

12. Francis S, Croft D, Schuttelkopf AW, Parry C, Pugliese A, Cameron K, Claydon S, Drysdale M, Gardner C, Gohlke A, Goodwin G, Gray CH, Konczal J, McDonald L, Mezna M, Pannifer A, Paul NR, Machesky L, McKinnon H, Bower J. Structure-based design, synthesis and biological evaluation of a novel series of isoquinolone and pyrazolo[4,3-c]pyridine inhibitors of fascin 1 as potential anti-metastatic agents. Bioorg Med Chem Lett. 2019;29(8):1023–9. Epub 2019/02/19. doi: 10.1016/j.bmcl.2019.01.035. PubMed PMID: 30773430; PMCID: PMC6419574.

13. Huang FK, Han S, Xing B, Huang J, Liu B, Bordeleau F, Reinhart-King CA, Zhang JJ, Huang XY. Targeted inhibition of fascin function blocks tumour invasion and metastatic colonization. Nature communications. 2015;6:7465. doi: 10.1038/ncomms8465. PubMed PMID: 26081695.

14. Yamakita Y, Matsumura F, Yamashiro S. Fascin1 is dispensable for mouse development but is favorable for neonatal survival. Cell motility and the cytoskeleton. 2009;66(8):524–34. Epub 2009/04/04. doi: 10.1002/cm.20356. PubMed PMID: 19343791; PMCID: 2738588.

15. Chen L, Yang S, Jakoncic J, Zhang JJ, Huang XY. Migrastatin analogues target fascin to block tumour metastasis. Nature. 2010;464(7291):1062–6. Epub 2010/04/16. doi: 10.1038/nature08978. PubMed PMID: 20393565; PMCID: 2857318.

16. Huang F-K, Han S, Xing B, Huang J, Liu B, Bordeleau F, Reinhart-King CA, Zhang JJ, Huang X-Y. Targeted inhibition of fascin function blocks tumour invasion and metastatic colonization. Nature communications. 2015;6:7465. doi: 10.1038/ncomms8465 https://www.nature.com/articles/ncomms8465#supplementary-information.

17. Alburquerque-Gonzalez B, Bernabe-Garcia A, Bernabe-Garcia M, Ruiz-Sanz J, Lopez-Calderon FF, Gonnelli L, Banci L, Pena-Garcia J, Luque I, Nicolas FJ, Cayuela-Fuentes ML, Luchinat E, Perez-Sanchez H, Montoro-Garcia S, Conesa-Zamora P. The FDA-Approved Antiviral Raltegravir Inhibits Fascin1-Dependent Invasion of Colorectal Tumor Cells In Vitro and In Vivo. Cancers (Basel). 2021;13(4). Epub 2021/03/07. doi: 10.3390/cancers13040861. PubMed PMID: 33670655.

18. Schild T, Low V, Blenis J, Gomes AP. Unique Metabolic Adaptations Dictate Distal Organ-Specific Metastatic Colonization. Cancer Cell. 2018;33(3):347–54. Epub 2018/03/14. doi: 10.1016/j.ccell.2018.02.001. PubMed PMID: 29533780; PMCID: PMC5889305.

19. Lunt SY, Vander Heiden MG. Aerobic glycolysis: meeting the metabolic requirements of cell proliferation. Annu Rev Cell Dev Biol. 2011;27:441–64. doi: 10.1146/annurev-cellbio-092910-154237. PubMed PMID: 21985671.

20. Hu H, Juvekar A, Lyssiotis CA, Lien EC, Albeck JG, Oh D, Varma G, Hung YP, Ullas S, Lauring J, Seth P, Lundquist MR, Tolan DR, Grant AK, Needleman DJ, Asara JM, Cantley LC, Wulf GM. Phosphoinositide 3-Kinase Regulates Glycolysis through Mobilization of Aldolase from the Actin Cytoskeleton. Cell. 2016;164(3):433–46. Epub 2016/01/30. doi: 10.1016/j.cell.2015.12.042. PubMed PMID: 26824656; PMCID: PMC4898774.

21. Elgendy M, Ciro M, Hosseini A, Weiszmann J, Mazzarella L, Ferrari E, Cazzoli R, Curigliano G, DeCensi A, Bonanni B, Budillon A, Pelicci PG, Janssens V, Ogris M, Baccarini M, Lanfrancone L, Weckwerth W, Foiani M, Minucci S. Combination of Hypoglycemia and Metformin Impairs Tumor Metabolic Plasticity and Growth by Modulating the PP2A-GSK3beta-MCL-1 Axis. Cancer Cell. 2019;35(5):798–815 e5. Epub 2019/04/30. doi: 10.1016/j.ccell.2019.03.007. PubMed PMID: 31031016.

22. Vander Heiden MG, Cantley LC, Thompson CB. Understanding the Warburg effect: the metabolic requirements of cell proliferation. Science. 2009;324(5930):1029–33. doi: 10.1126/science.1160809. PubMed PMID: 19460998; PMCID: 2849637.

23. Yang S, Huang FK, Huang J, Chen S, Jakoncic J, Leo-Macias A, Diaz-Avalos R, Chen L, Zhang JJ, Huang XY. Molecular mechanism of fascin function in filopodial formation. J Biol Chem. 2013;288(1):274–84. Epub 2012/11/28. doi: 10.1074/jbc.M112.427971. PubMed PMID: 23184945; PMCID: 3537022.

24. Korobova F, Ramabhadran V, Higgs HN. An actin-dependent step in mitochondrial fission mediated by the ER-associated formin INF2. Science. 2013;339(6118):464–7. doi: 10.1126/science.1228360. PubMed PMID: 23349293; PMCID: 3843506.

25. Ashizawa K, Willingham MC, Liang CM, Cheng SY. In vivo regulation of monomer-tetramer conversion of pyruvate kinase subtype M2 by glucose is mediated via fructose 1,6-bisphosphate. J Biol Chem. 1991;266(25):16842–6. Epub 1991/09/05. PubMed PMID: 1885610.

26. Park JS, Burckhardt CJ, Lazcano R, Solis LM, Isogai T, Li L, Chen CS, Gao B, Minna JD, Bachoo R, DeBerardinis RJ, Danuser G. Mechanical regulation of glycolysis via cytoskeleton architecture. Nature. 2020;578(7796):621–6. Epub 2020/02/14. doi: 10.1038/s41586-020-1998-1. PubMed PMID: 32051585; PMCID: PMC7210009.

27. Koo JH, Guan KL. Interplay between YAP/TAZ and Metabolism. Cell Metab. 2018;28(2):196–206. Epub 2018/08/09. doi: 10.1016/j.cmet.2018.07.010. PubMed PMID: 30089241.

28. Dupont S, Morsut L, Aragona M, Enzo E, Giulitti S, Cordenonsi M, Zanconato F, Le Digabel J, Forcato M, Bicciato S, Elvassore N, Piccolo S. Role of YAP/TAZ in mechanotransduction. Nature. 2011;474(7350):179–83. doi: 10.1038/nature10137. PubMed PMID: 21654799.

29. Fornes O, Castro-Mondragon JA, Khan A, van der Lee R, Zhang X, Richmond PA, Modi BP, Correard S, Gheorghe M, Baranasic D, Santana-Garcia W, Tan G, Cheneby J, Ballester B, Parcy F, Sandelin A, Lenhard B, Wasserman WW, Mathelier A. JASPAR 2020: update of the open-access database of transcription factor binding profiles. Nucleic acids research. 2020;48(D1):D87–D92. Epub 2019/11/09. doi: 10.1093/nar/gkz1001. PubMed PMID: 31701148; PMCID: PMC7145627.

30. Lin S, Huang C, Sun J, Bollt O, Wang X, Martine E, Kang J, Taylor MD, Fang B, Singh PK, Koomen J, Hao J, Yang S. The mitochondrial deoxyguanosine kinase is required for cancer cell stemness in lung adenocarcinoma. EMBO molecular medicine. 2019;11(12):e10849. Epub 2019/10/22. doi: 10.15252/emmm.201910849. PubMed PMID: 31633874; PMCID: PMC6895611.

31. Pelosi G, Pastorino U, Pasini F, Maissoneuve P, Fraggetta F, Iannucci A, Sonzogni A, De Manzoni G, Terzi A, Durante E, Bresaola E, Pezzella F, Viale G. Independent prognostic value of fascin immunoreactivity in stage I nonsmall cell lung cancer. British journal of cancer. 2003;88(4):537–47. Epub 2003/02/20. doi: 10.1038/sj.bjc.6600731. PubMed PMID: 12592367; PMCID: 2377175.

32. Tiriac H, Belleau P, Engle DD, Plenker D, Deschenes A, Somerville TDD, Froeling FEM, Burkhart RA, Denroche RE, Jang GH, Miyabayashi K, Young CM, Patel H, Ma M, LaComb JF, Palmaira RLD, Javed AA, Huynh JC, Johnson M, Arora K, Robine N, Shah M, Sanghvi R, Goetz AB, Lowder CY, Martello L, Driehuis E, LeComte N, Askan G, Iacobuzio-Donahue CA, Clevers H, Wood LD, Hruban RH, Thompson E, Aguirre AJ, Wolpin BM, Sasson A, Kim J, Wu M, Bucobo JC, Allen P, Sejpal DV, Nealon W, Sullivan JD, Winter JM, Gimotty PA, Grem JL, DiMaio DJ, Buscaglia JM, Grandgenett PM, Brody JR, Hollingsworth MA, O’Kane GM, Notta F, Kim E, Crawford JM, Devoe C, Ocean A, Wolfgang CL, Yu KH, Li E, Vakoc CR, Hubert B, Fischer SE, Wilson JM, Moffitt R, Knox J, Krasnitz A, Gallinger S, Tuveson DA. Organoid Profiling Identifies Common Responders to Chemotherapy in Pancreatic Cancer. Cancer Discov. 2018;8(9):1112–29. Epub 2018/06/02. doi: 10.1158/2159-8290.CD-18-0349. PubMed PMID: 29853643; PMCID: PMC6125219.

33. Jackson EL, Olive KP, Tuveson DA, Bronson R, Crowley D, Brown M, Jacks T. The differential effects of mutant p53 alleles on advanced murine lung cancer. Cancer Res. 2005;65(22):10280–8. Epub 2005/11/17. doi: 10.1158/0008-5472.CAN-05-2193. PubMed PMID: 16288016.

34. Li A, Morton JP, Ma Y, Karim SA, Zhou Y, Faller WJ, Woodham EF, Morris HT, Stevenson RP, Juin A, Jamieson NB, MacKay CJ, Carter CR, Leung HY, Yamashiro S, Blyth K, Sansom OJ, Machesky LM. Fascin is regulated by slug, promotes progression of pancreatic cancer in mice, and is associated with patient outcomes. Gastroenterology. 2014;146(5):1386–96 e1-17. Epub 2014/01/28. doi: 10.1053/j.gastro.2014.01.046. PubMed PMID: 24462734; PMCID: 4000441.

35. Zhao X, Gao S, Ren H, Sun W, Zhang H, Sun J, Yang S, Hao J. Hypoxia-inducible factor-1 promotes pancreatic ductal adenocarcinoma invasion and metastasis by activating transcription of the actin-bundling protein fascin. Cancer Res. 2014;74(9):2455–64. Epub 2014/03/07. doi: 10.1158/0008-5472.CAN-13-3009. PubMed PMID: 24599125.

36. Bernstein BW, Bamburg JR. Actin-ATP hydrolysis is a major energy drain for neurons. J Neurosci. 2003;23(1):1–6. Epub 2003/01/07. PubMed PMID: 12514193; PMCID: PMC6742122.

37. Masters C. Interactions between glycolytic enzymes and components of the cytomatrix. J Cell Biol. 1984;99(1 Pt 2):222s–5s. Epub 1984/07/01. doi: 10.1083/jcb.99.1.222s. PubMed PMID: 6746730; PMCID: PMC2275576.

38. Kang J, Wang J, Yao Z, Hu Y, Ma S, Fan Q, Gao F, Sun Y, Sun J. Fascin induces melanoma tumorigenesis and stemness through regulating the Hippo pathway. Cell Commun Signal. 2018;16(1):37. Epub 2018/07/05. doi: 10.1186/s12964-018-0250-1. PubMed PMID: 29970086; PMCID: PMC6029074.

39. Liang Z, Wang Y, Shen Z, Teng X, Li X, Li C, Wu W, Zhou Z, Wang Z. Fascin 1 promoted the growth and migration of non-small cell lung cancer cells by activating YAP/TEAD signaling. Tumour Biol. 2016;37(8):10909–15. doi: 10.1007/s13277-016-4934-0. PubMed PMID: 26886283.

40. DuPage M, Dooley AL, Jacks T. Conditional mouse lung cancer models using adenoviral or lentiviral delivery of Cre recombinase. Nature protocols. 2009;4(7):1064–72. doi: 10.1038/nprot.2009.95. PubMed PMID: 19561589; PMCID: PMC2757265.

41. Yang S, Zhang JJ, Huang XY. Mouse models for tumor metastasis. Methods Mol Biol. 2012;928:221–8. Epub 2012/09/08. doi: 10.1007/978-1-62703-008-3_17. PubMed PMID: 22956145.

42. Sun J, Lu F, He H, Shen J, Messina J, Mathew R, Wang D, Sarnaik AA, Chang WC, Kim M, Cheng H, Yang S. STIM1- and Orai1-mediated Ca2+ oscillation orchestrates invadopodium formation and melanoma invasion. J Cell Biol. 2014;207(4):535–48. Epub 2014/11/19. doi: 10.1083/jcb.201407082. PubMed PMID: 25404747; PMCID: 4242838.

43. Tang Z, Li C, Kang B, Gao G, Li C, Zhang Z. GEPIA: a web server for cancer and normal gene expression profiling and interactive analyses. Nucleic acids research. 2017;45(W1):W98–W102. Epub 2017/04/14. doi: 10.1093/nar/gkx247. PubMed PMID: 28407145; PMCID: PMC5570223.

44. Gunda V, Yu F, Singh PK. Validation of metabolic alterations in microscale cell culture lysates using hydrophilic interaction liquid chromatography (HILIC)-tandem mass spectrometry-based metabolomics. PloS one. 2016;11(4).

45. Yuan M, Breitkopf SB, Yang X, Asara JM. A positive/negative ion–switching, targeted mass spectrometry–based metabolomics platform for bodily fluids, cells, and fresh and fixed tissue. Nature protocols. 2012;7(5):872.

46. Trefely S, Ashwell P, Snyder NW. FluxFix: automatic isotopologue normalization for metabolic tracer analysis. BMC Bioinformatics. 2016;17(1):485. doi: 10.1186/s12859-016-1360-7. PubMed PMID: 27887574; PMCID: PMC5123363.

